# Sensory processing reformats odor coding around valence and dynamics

**DOI:** 10.1101/2025.11.08.687380

**Authors:** Kristyn M. Lizbinski, Kay J Ellison, Gizem Sancer, Helen X. Mao, James M. Jeanne

## Abstract

Extracting relevant features of a complex sensory signal typically involves sequential processing through multiple brain regions. However, identifying the logic and mechanisms of these transformations has been difficult, due to the challenges of measuring both activity within and long-range connectivity between multiple neural populations. Here, we investigate the reformatting of odor information across two stages of the *Drosophila* olfactory system. We measure the odor tuning of 20 types of anatomically-defined third order lateral horn neuron (LHN) and compare to predictions based on the odor tuning of second-order projection neurons (PNs) and PN-LHN connectivity. We find that LHNs reformat PN activity in two distinct ways. First, LHNs selectively discard information about odor identities with similar valence (i.e., attractiveness or aversiveness). This emerges from a precise alignment of PN odor tuning and PN-LHN connectivity, as well as odor-specific inhibition and boosting of LHN activity. This creates a population code for valence that is more explicit than in PNs. Second, a subset of LHNs selectively discard information about continuing odor presence, by responding only transiently to odor onset. This creates a population code for odor dynamics that is more explicit than in PNs. Across LHNs, valence and dynamics are independent of each other. Thus, feedforward connectivity and local inhibition combine to extract two orthogonal dimensions of olfactory information.

## INTRODUCTION

Perceptual judgements typically rely on the precise curation of vast amounts of sensory information. This requires parallel processing by large neural populations as well as sequential processing by hierarchically interconnected brain regions. Current deep learning models that link parallel and sequential processing to perception lack mechanistic precision^1^, meaning that they do not account for true interactions between neural coding and connectivity. Identifying these interactions remains challenging because it requires characterization of the neural code of complete (or nearly complete) neural populations, along with a comprehensive mapping of connections between neurons in different brain regions. Consequently, an understanding of how feedforward circuits transform implicit neural codes into more explicit ones has remained elusive.

The specific pattern of feedforward connectivity between hierarchical layers is a major determinant of sensory processing^2,3^. Because interconnected brain regions are often spatially distant, complete mapping of inter-regional connectivity at single synapse resolution in vertebrates is still beyond reach. However, the brain of the fruit fly, *Drosophila*, is small enough that connectomes now include all long-distance projections^4–6^. This means that neural coding in multiple regions can be directly linked to their exact interconnectivity.

Here, we take advantage of the compactness of the *Drosophila* olfactory system to relate a nearly complete population code and its exact pattern of downstream connections to the formation of an explicit neural code for innate odor valence. As in vertebrate olfaction, odors bind to receptors housed in olfactory receptor neurons (ORNs)^7^. Each ORN typically expresses a single receptor type, which defines its receptive field^8^, and all ORNs that express the same receptor project their axons into the same glomerulus. The full array of 51 glomeruli thus forms the complete olfactory information channel^9–11^. PNs carry glomerular output to target regions, where they converge and diverge onto LHNs^12,13^, which collectively encode the innate valence of odors^14^, and drive attraction or aversion behavior^15,16^.

In this study, we investigate how the PN population code is transformed by its pattern of axonal projections onto LHNs. We find that PN odor coding aligns with feedforward PN-LHN connectivity to create anatomically distinct channels for attractive and aversive valence in the lateral horn. The PN odor code – combined with feedforward connectivity – accurately predicts the responses of many LHNs. However, some LHNs operate under strong odor-specific inhibition, which suppresses responses to attractive odors. Other LHNs rely on odor-specific integration, which boosts responses to attractive odors. Finally, both attractive and aversive LHNs can have either transient or sustained odor responses, indicating a broad bifurcation of temporal dynamics across the LHN population that spans the full spectrum of valence. Together, this reveals how sequential processing stages transform an implicit and temporally homogeneous valence code into a more explicit and temporally diverse format. More generally, it provides a framework for investigating computation by linking neural coding with connectivity.

## RESULTS

### The lateral horn encodes odor valence more explicitly than the antennal lobe

We began by measuring the innate behavioral valence for each of 12 odors in a two-choice assay. Flies walked freely in an arena, while odor was delivered to one side and clean air to the other (**Figure 1A**). The 12 odors had a range of valences (quantified as preference index; **Figure 1B**; Methods), with benzaldehyde the most negative and apple cider vinegar (ACV) the most positive, as expected^17–19^. To understand how the brain forms explicit representations of valence, we first compared the PN odor code to the LHN odor code.

**Figure 1.**
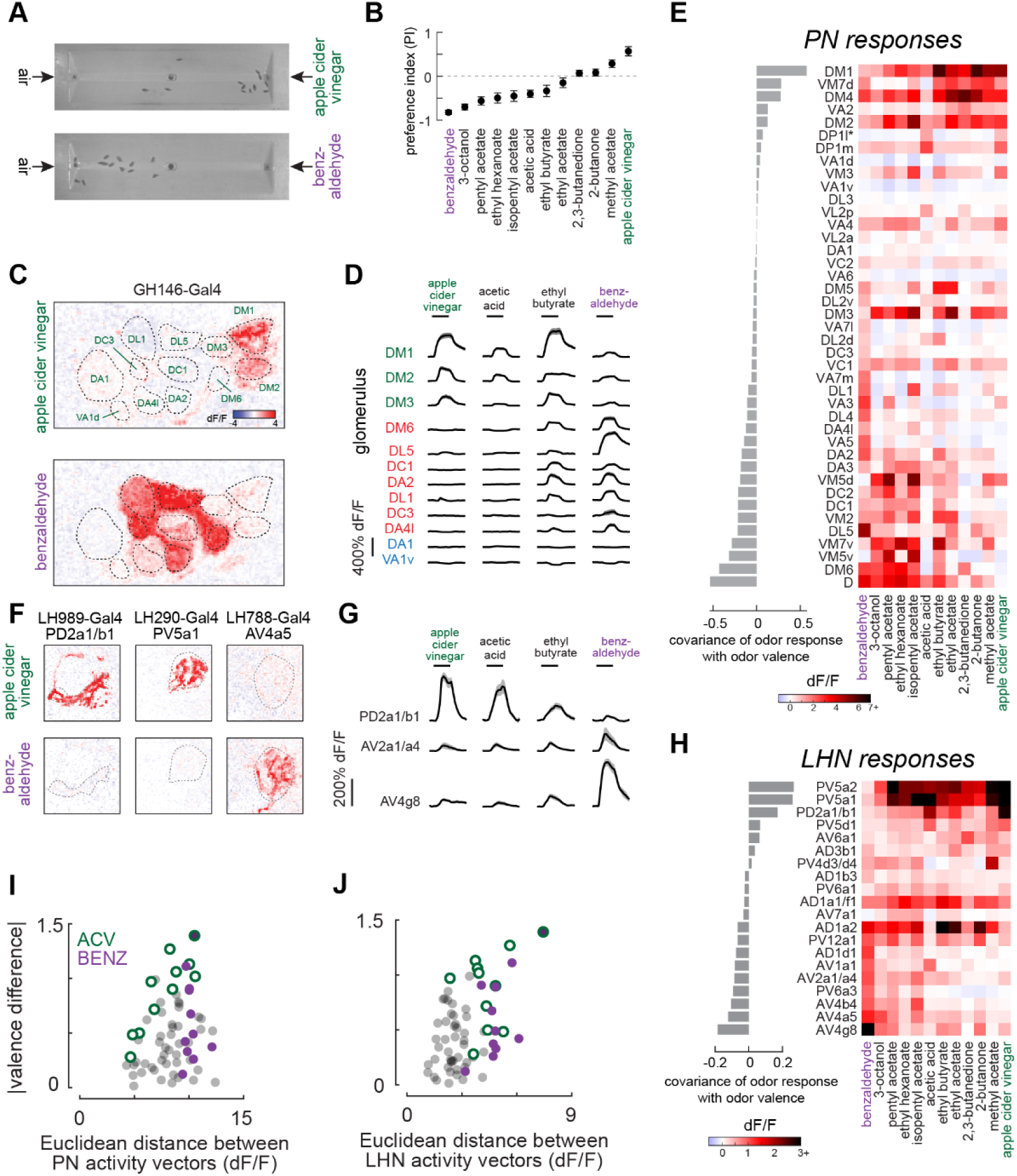
LHNs encode valence better than PNs. **(A)** Example still images from behavioral valence place-preference assay. Flies prefer ACV compared to clean air and prefer clean air to benzaldehyde. **(B)** Mean (± s.e.m.) preference indices for each of 12 odors (all tested against clean air). Preference index (PI) was calculated at each time point as (*NN_o_*_dor_ – *NN_a_*_ir_)/*NN_t_*_otal_, where *NN_o_*_dor_ and *NN_a_*_ir_ are the numbers of flies in the halves of the arena closer to the odor and air inlets, respectively. Each point is the average of 5-7 cohorts, with 10-12 flies per cohort. **(C)** Odor responses in a representative plane of the antennal lobe measured with *in vivo* 2-photon imaging of GCaMP6f fluorescence. The glomerulus boundaries and identities are identical for both odors, because they are the same plane of the same fly. **(D)** Mean (± s.e.m.) fluorescence responses (dF/F) across flies (n = 5-24, except for DC1 responses to benzaldehyde, which is n = 3 flies) to 4 odors for each of the glomeruli identified in (C). All measurements are from cholinergic uniglomerular PNs, except for glomerulus DP1l (marked with an asterisk) where ORNs were measured. **(E)** (Right) Heatmap of mean responses across flies for each odor and each glomerulus (n = 2-24, median sample size = 10 flies). (Left) covariance of odor responses with preference index for each glomerulus. **(F)** Responses to two odors in three representative LHN types measured with *in vivo* 2-photon imaging, as in (C). **(G)** Same as (D) but for representative LHN types (n = 11, 5, 5 flies for PD2a1/b1, AV2a1/a4, AV4g8, respectively). **(H)** Same as (E) but for LHNs (average of 3-19 flies per LHN type, median sample size = 5 flies). **(I)** Relationship between Euclidean distances between PN activity vectors and the absolute value of the valence difference for each of the 66 odor pairs. Pearson correlation, r = 0.27, p = 0.03). **(J)** Relationship between Euclidean distances between LHN activity vectors and the absolute value of the valence difference for each of the 66 odor pairs. Pearson correlation, r = 0.48, p = 4.7×10^-5^). This relationship is stronger than the corresponding relationship for PNs (panel I). ANCOVA interaction term, p = 0.026).

We measured the responses of PNs innervating 40 of the 51 olfactory glomeruli to the same odors using GCaMP6f under the control of either GH146-Gal4^20^ or VT033006-Gal4^21^ (**Figure 1C-E**). The remaining glomeruli were omitted because they are either not innervated by canonical uniglomerular PNs, were not easily identifiable within the antennal lobe, or no known driver line exists to label them. The one exception was glomerulus DP1l, from which we imaged from cognate ORNs instead of PNs. Thus, we consider the activity of 41 glomeruli. Using identical odor delivery, we then measured LHN activity using GCaMP6f under the control of 20 selective split-Gal4 lines^16,22^ (**Figure 1F-H, Table S1**). Most of these lines label single LHN types, but a few label multiple types (we also determined that one line, LH788-Gal4, labels AV4a5 LHNs, which is different than previously reported^16^; **Figure S1**).

As expected, certain PNs organized their odor responses by valence^23^. PNs innervating glomeruli driving attraction (e.g., e.g., DM1 and VA2^17,24^) had positive covariances between neural and behavioral responses. PNs that innervate glomeruli driving aversion (DL5 and DA2^25,26^) had negative covariances between neural and behavioral responses (**Figure 1E**).

Certain LHNs also organized their odor responses by valence. LHN types that drive attraction (e.g., PD2a1/b1 and PV5a1^15,16,27^) had positive covariances between neural and behavioral responses. LHN types that drive aversion (e.g., AV1a1^16,28^) had negative covariances between neural and behavioral responses (**Figure 1H**). Although our sample of LHN types was less comprehensive than our sample of PNs, the range of valence coding we observed was statistically indistinguishable from a larger sample measured from LHN somata (which precluded identification of LHN types; **Figure S2**)^14^. Thus, the subset of LHNs we investigate here are a representative sample of the whole population, in terms of valence coding.

While some individual PNs and LHNs show strong relationships to valence, odor coding generally depends on larger populations^8^. We thus compared how well PN and LHN populations encode valence based on their odor tuning. To consider each sampled population fully, we considered each neuron’s response as a dimension in neural activity space and assembled activity vectors for each odor for PNs and LHNs^29,30^. Valence encoding can then be assessed by relating Euclidean distances between activity vectors for pairs of odors to the corresponding difference in odor valence. PNs exhibited no relationship between vector distance and valence difference across all odor pairs (**Figure 1I**). This occurred because many odor pairs are highly distinguishable by PNs but nonetheless have similar innate valences (i.e., points at the lower right in **Figure 1I**). In contrast, LHNs exhibited a consistent relationship between vector distance and valence (**Figure 1J**). This indicates that the LHN population encodes valence more explicitly than the PN population (in agreement with recent findings^14^), at least in part by sacrificing the ability to discriminate between odors of similar valence.

### PN-LHN connectivity compresses representations of similar valence

Most LHNs receive convergent input from multiple PN types^12,13^. Convergence discards information (since the identity of each input is lost), so the specific pattern of which PN types converge will constrain the information available to LHNs. Acknowledging that other circuit components and nonlinear processes may provide further refinement, we began by hypothesizing that PNs with similar odor coding valence tend to converge onto the same LHNs, while LHNs of different valence avoid convergence onto the same LHNs.

To test this hypothesis, we compared PN odor coding valences (defined as the covariance between neural and behavioral responses, **Figure 1E**) with anatomic PN-LHN connection strength (defined as the number of synapses), for each LHN in the Hemibrain connectome^4^. Some LHNs receive input from PNs with similar odor coding valences (e.g., **Figure 2A**). When we shuffled the connectivity (but preserved the total number of output synapses from each PN), this tendency diminished, meaning that LHNs are more likely to receive inputs with similar valence coding than expected by chance (**Figure 2B**).

**Figure 2.**
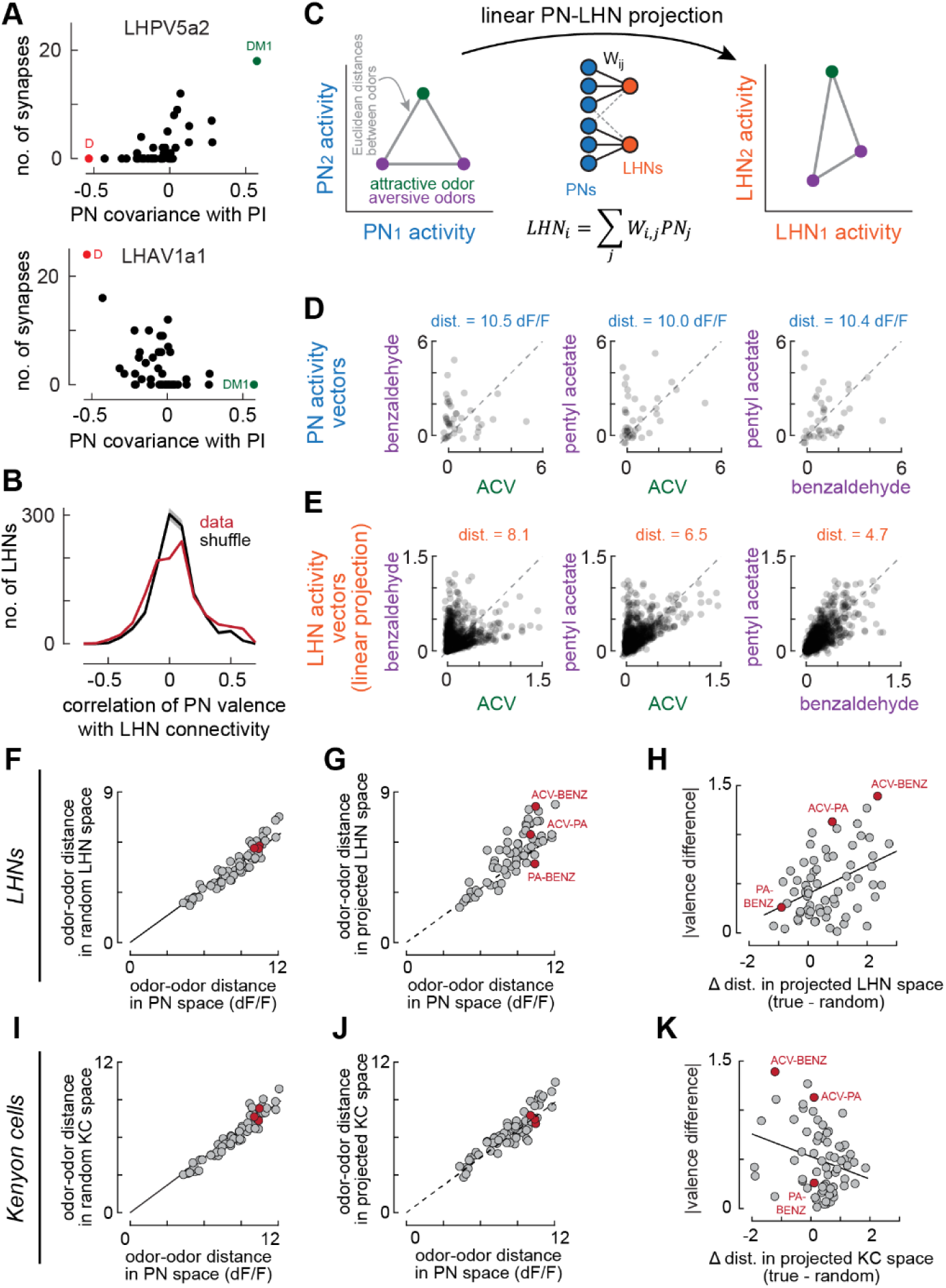
PN-LHN connectivity compresses representations of similar valence. **(A)** Relationship between PN odor coding valence (i.e., PN response covariance with PI, reported in Figure 1E) and PN connection weights for two example LHNs. PN weights for the example PV5a2 LHN (top) are positively correlated with odor coding valence (Pearson correlation, r = 0.67). PN weights for the example AV1a1 LHN (bottom) are negatively correlated with odor coding valence (Pearson correlation, r = −0.53). **(B)** Distribution of Pearson correlations, as in (A), for all 1476 LHNs in the Hemibrain connectome (red). Shuffling PN-LHN connectivity reduced the number of LHNs receiving PN inputs with similar odor coding valence. Kolmogorov-Smirnov test, p = 4.2×10^-4^. **(C)** 2-D schematic of activity vectors and the procedure for linear projection of PN activity into LHN space. The linear projection is defined by PN-LHN connection weights in the Hemibrain connectome. **(D)** PN activity vectors between all pairs of three odors. Each point is the average response of a single PN. The Euclidean distance between each pair of vectors is shown at the top of each plot. **(E)** Same as (D), but for projected LHN activity vectors. **(F)** Euclidean distances between PN activity vectors for all pairs of odors compared to Euclidean distances between projected LHN activity vectors built from shuffled PN-LHN connectivity. Solid line is a linear regression constrained to pass through the origin. Red points identify the same odor pairs shown in (D,E). **(G)** Euclidean distances between PN activity vectors for all pairs of odors compared to Euclidean distances between LHN activity vectors built from actual PN-LHN connectivity. Dashed line is the same linear regression from (F), highlighting deviations from the shuffle control. **(H)** Deviations of LHN distances from the shuffle control (F,G) increase with valence differences (Pearson correlation, r = 0.42, p = 0.0005). **(I,J)** Same as (F,G) but for 1828-dimensional KC space instead of LHN space. Euclidian distances between projected KC activity vectors are more similar to the shuffled control than for LHNs. **(K)** Same as (H) but for KCs. Deviations of KC distances from the shuffle control modestly decrease with valence differences (Pearson correlation, r = −0.27, p = 0.03).

This pattern of connectivity could, in principle, compress the representations of odors with similar valence (i.e., by blurring the distinction between them in LHN activity space). To investigate this, we applied the same population vector distance analysis as above, except we now modeled the responses of every LHN as a weighted linear sum of PN responses, with the weights determined by the Hemibrain connectome (we refer to this as the linear PN-LHN projection, Methods; **Figure 2C**). This allowed us to focus exclusively on the role of PN-LHN connectivity, as it pertains to the odors in our panel. To illustrate this approach, we consider the population vectors in both PN and LHN activity space for three odors (**Figure 2D,E**). PN vectors were equivalently dissimilar for all three odors (due to many points being far from the unity line for each odor pair in **Figure 2D**). LHN vectors, in contrast, were more variable for these odors (due to distinct patterns around the unity line for different odor pairs in **Figure 2E**). PN-LHN connectivity therefore transforms representations that distinguish some odors, but not others.

Odor-odor distances in PN space and LHN space are not directly comparable, due to different numbers of neurons, and an unconstrained scaling factor on each synapse (i.e., we treat each connection weight as a relative value, not an absolute value). To make a fair comparison, we began by randomly shuffling the PN-LHN connectivity matrix. This preserves the total number of LHNs and the total distribution of connection weights but eliminates structured connectivity. Odor-odor distances in this random LHN space were linearly related to odor-odor distances in PN space (**Figure 2F**), consistent with random connectivity impairing representations of all odors equally. Certain odor-odor distances in the true projected LHN space, however, were larger than in the random space (**Figure 2G**). Most importantly, distances were greater for odor pairs with larger valence differences (**Figure 2H**). Thus, PN-LHN connectivity squeezes representations together for odors with similar valence more than it squeezes representations for odors with different valence.

In addition to LHNs, PNs also target Kenyon cells (KCs), the intrinsic neurons of the mushroom body, which forms learned odor associations. Because PN-KC connectivity is less structured^13,31–33^ than PN-LHN connectivity, we hypothesized that it does not align with PN valence. Surprisingly, the same odor-odor distance analysis in KC space revealed a modest negative relationship: PN-KC connectivity de-emphasizes valence (**Figure 2I-K**). Nevertheless, this demonstrates that the alignment of PN-LHN connectivity with valence is not a universal property of each PN, indicating a specialization of the lateral horn for innate valence.

### PN-LHN connectivity defines anatomical channels of opposing valence

The only way that the linear transformation implemented by the PN-LHN connectivity matrix can preserve representations of different valence, while diminishing representations of similar valence, is by maintaining segregated channels of connectivity. In other words, PNs encoding attractive odors must largely avoid the LHNs targeted by PNs encoding aversive odors, and vice versa. We thus analyzed PN-LHN convergence patterns to identify PNs and LHNs that belong to these segregated channels.

Without any priors on valence, we computed the pairwise correlation matrix between PN connectivity patterns. As expected^12,34^, hierarchical clustering revealed several broad channels of PN types (**Figure 3A**). One channel contained mostly pheromone responsive PNs which we did not explicitly target with our odor panel and thus excluded from further analysis. The remaining two channels corresponded to PNs with distinct axonal anatomy^34,35^. “Channel 1” PN axons targeted a dorsal “core” of the lateral horn, while “channel 2” PN axons either targeted a more ventral region, or else formed claws that wrapped around this core (**Figure 3B,C, S3**).

**Figure 3.**
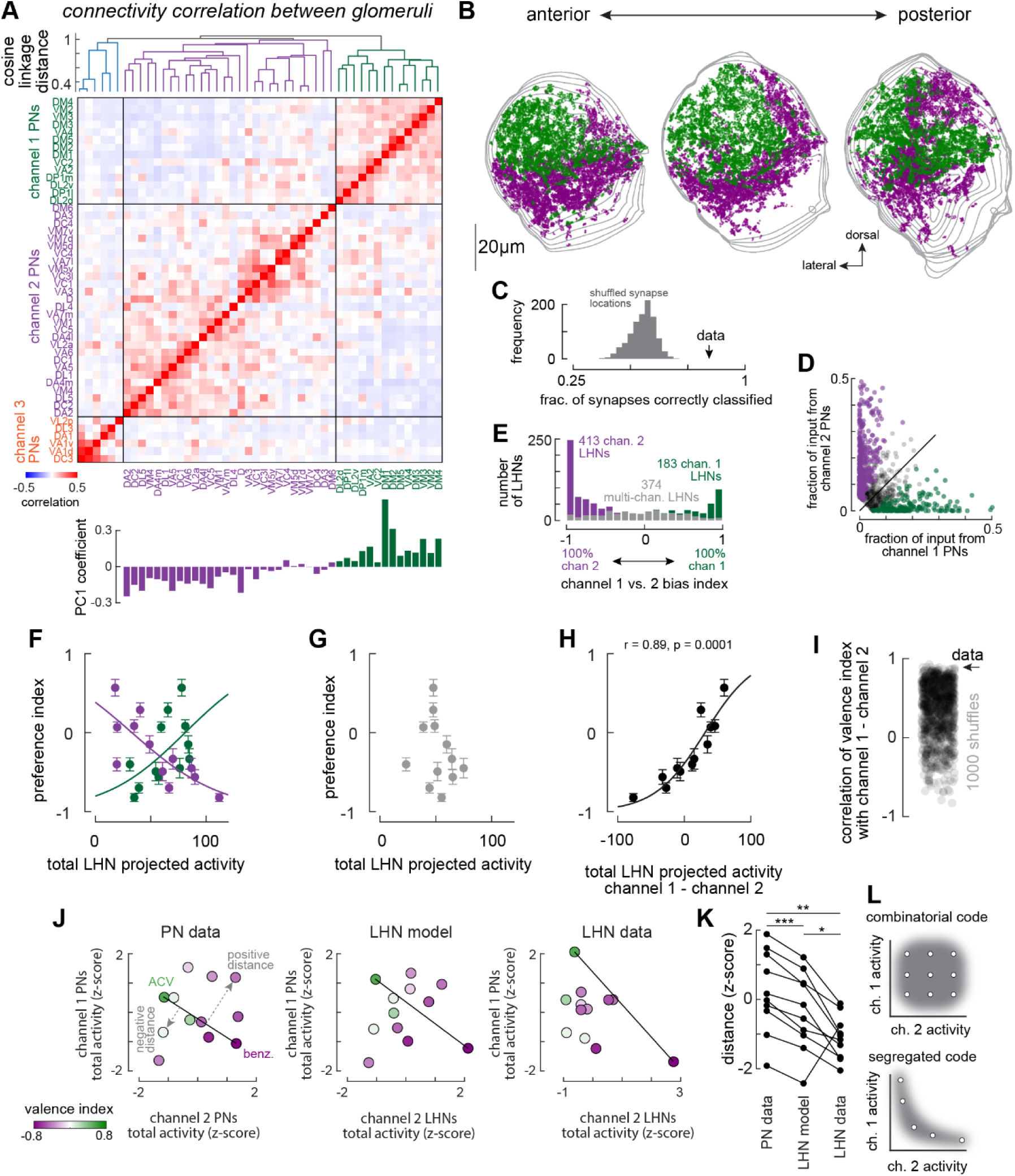
PN-LHN connectivity defines valence channels in the lateral horn and transform a combinatorial code into a segregated code. **(A)** Top: dendrogram of hierarchical clustering based on cosine distance between PN connectivity patterns with LHNs, which forms three broad channels. Middle: connectivity correlation between all pairs of PNs based on their anatomic connectivity with LHNs. Each value in the symmetric matrix is the correlation coefficient between a single pair of PN types. Bottom: coefficients of the first principal component of channel 1 and 2 PN connectivity separate the connectivity in a similar manner as the hierarchical clustering analysis. **(B)** Locations of all lateral horn presynapses and postsynapses from channel 1 (green, 19,550 synapses) and channel 2 (purple, 27,519 synapses) PNs. Each image is a projection into the dorsal-lateral plane of synapses locations occurring within a 36μm (left and right projections) or 12μm (center) “slab” of the lateral horn (gray contours are the boundaries of the LH at every 3μm through the slab. **(C)** Spatial segregation into two regions was quantified as the fraction of synapses on the correct side of a 2-D manifold embedded in 3-D space, defined by a support vector machine (black arrow). 1000 random shuffles of the channel membership of each PN (and recomputing the 2-D manifold) yielded a distribution of correct classifications significantly less than for the true channel membership. **(D)** Comparison of the fraction of input from channel 1 and channel 2 PNs for all 970 LHNs that receive at least 5% of their total input from channel 1 and 2 PNs. LHNs with biased inputs from channel 1 or channel 2 (*χχ^2^* test vs. Bernoulli distribution with equal probabilities, corrected for 970 comparisons; Methods) are colored green and purple, respectively. Unbiased (“multi-channel”) LHNs are grey. **(E)** Distribution of channel 1 vs. 2 bias index (difference between channel 1 and 2 PN inputs normalized by their sum) for all LHNs in (C). 61% of LHNs are significantly biased for either channel 1 or channel 2 inputs. **(F)** Total projected odor responses for all channel 1 LHNs (green) and channel 2 LHNs (purple) correspond to odor valence (Pearson correlation: channel 1, r = −0.77, p = 0.0035; channel 2, r = 0.58, p = 0.049). Curves are sigmoid fits. **(G)** Total projected odor responses for all multi-channel LHNs do not correspond to odor valence (Pearson correlation: r = −0.22, p = 0.49). **(H)** The difference between projected channel 1 and channel 2 activity correlates strongly with valence (r = 0.89, p = 0.0001). Curve is a sigmoid fit. **(I)** Independently shuffling each PN’s output connectivity (preserving total synaptic output from each PN) impairs valence coding (quantified as the difference between projected channel 1 and channel 2 total activity). **(J)** Comparison of total channel 1 and channel 2 activity for each of the 12 odors, for PN activity data (left), projected LHN activity (simulated from PN activity data and PN-LHN connectivity; center), and for LHN activity data (right). The projected LHN activity only includes the LHN types for which actual responses were measured. These 2-dimensional distributions were characterized by measuring the distance from each odor to the “ACV-benzaldehyde” line (i.e., the axis linking the two strongest valences; see K,L). **(K)** Distances to the ACV-benzaldehyde line are more negative for projected LHN activity than PN activity, and more negative for true LHN activity than projected LHN activity. T-tests: * p = 0.039; ** p = 0.0029; *** p = 1.3×10^-5^. **(L)** PN activity resembles a combinatorial code (top), where different odors fully span the space of possible combinations of channel 1 and channel 2 activity. LHN activity resembles a segregated code, where different odors activate either channel alone, but not both channels simultaneously.

We then quantified the fraction of input from channel 1 and channel 2 PNs for each LHN in the connectome. We identified 183 and 413 LHNs with significantly biased input from channel 1 or 2 PNs, respectively (**Figure 3D,E**). These two LHN channels also formed anatomically distinct projection patterns, with channel 1 biased towards the superior intermediate protocerebrum and channel 2 biased towards the superior lateral protocerebrum (**Figure S3**).

Predicted odor responses (based on the linear transformation defined by PN-LHN connectivity) for channel 1 and 2 LHNs encoded positive and negative valence, respectively (**Figure 3F**). The difference between channel 1 and channel 2 LHN activity encoded valence even more accurately (**Figure 3G**). Randomly shuffling odor responses between PNs abolished valence coding in the simulated LHN activity (**Figure 3H,I**), meaning that PN odor coding and PN-LHN connectivity are coordinated around valence.

### LHNs transform the combinatorial PN code into a segregated code

Because channels of negative and positive odor valence remain largely segregated, they form a 2-dimensional coding space that is largely preserved between PNs and LHNs. This provides an equal footing to compare odor coding between these layers of processing. Our odor panel elicited activity that filled the coding space spanned by channel 1 and channel 2 PNs (**Figure 3J**), in line with the so-called combinatorial odor code across glomeruli^8,23^. ACV, the most attractive odor in our panel, evoked strong responses in channel 1 PNs.

Benzaldehyde, the most aversive odor in our panel, evoked strong responses in channel 2 PNs. However, multiple other odors evoked greater total PN activity than ACV and benzaldehyde did (i.e., points above the line connecting them in this PN space; **Figure 3J,K**). Therefore, neither PN channel encodes valence monotonically.

We then considered the linear model predictions for channel 1 and 2 LHNs (restricted to those types that we sampled experimentally). The linear transformation imposed by connectivity made ACV and benzaldehyde more prominent (fewer points above the line connecting them) than in PN space (**Figure 3J,K**). This indicates that connectivity contributes to a reformatting of the combinatorial PN code into a more segregated LHN code (**Figure 3L**).

Finally, we investigated actual odor-evoked activity in channel 1 and 2 LHNs. Compared to the linear model, real LHNs encoded ACV and benzaldehyde even more prominently (no odors evoked responses greater than their joint activity; **Figure 3J,K**), resembling a segregated code more than a combinatorial code (**Figure 3L**). This means that additional processing, beyond that imposed by connectivity, makes important contributions to a segregated valence code in LHNs.

### Asymmetric local inhibition reshapes odor coding

To further investigate the additional processing that contributes to LHN odor coding, we directly compared predictions of the feedforward linear model to actual odor responses in each sampled LHN type within the ‘aversive’ and ‘appetitive’ channels (i.e., channel 1 and channel 2, respectively). While the linear model predicted the overall landscape of odor responses in both channels, it performed better for aversive channel LHNs (**Figure 4A,B**).

**Figure 4.**
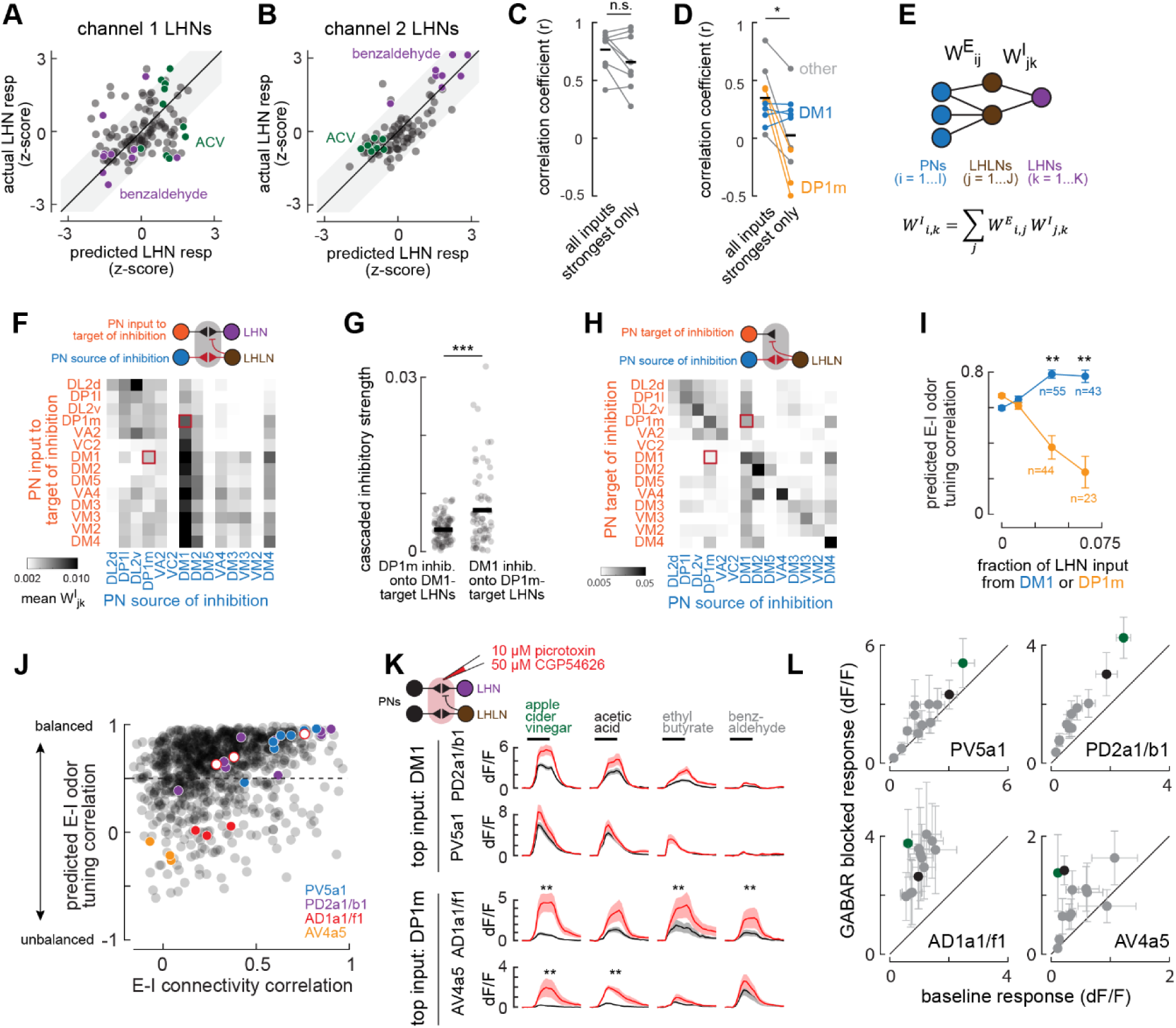
Asymmetric local inhibition reshapes odor coding. **(A)** Actual LHN responses vs. predicted responses from the feedforward linear model for appetitive (channel 1) LHNs. Each point is the average response of one LHN to one odor. Odor responses are z-scored within each LHN type. Pearson correlation: r = 0.35, p = 0.0001. Each LHN’s response to ACV and benzaldehyde are identified in green and purple, respectively. **(B)** Same as (A), but for aversive (channel 2) LHNs. Pearson correlation: r = 0.78, p = 7×10^-21^. **(C)** Accuracy of predictive models of odor coding for each aversive LHN type, quantified by Pearson correlation coefficient. **(D)** Same as (C), but for appetitive LHN odor coding. * t-test, p = 0.018. LHN types receiving strongest input from DM1 and DP1m PNs are highlighted. **(E)** Schematic of cascaded linear model to predict the inhibitory effects of each PN on each LHN, via interposed LHLNs. **(F)** Heatmap of cascaded PN-LHLN-LHN connection weights. Each row denotes the average cascaded weight for all LHNs with at least 2.5% of their total synaptic input from direct excitation of the specified PN. Columns denote the PN sources of inhibition. Red outlines highlight asymmetry between DP1m and DM1. **(G)** Quantification of asymmetry of inhibition between DM1 and DP1m onto target LHNs. Anatomical strength of inhibition arising from DM1 onto DP1m-target LHNs is stronger than inhibition arising from DP1m onto DM1target LHNs. *** t-test: p = 5.9×10^-6^. **(H)** Same as (F), except for PN-LHLN-PN connection weights. Each row denotes the average cascaded weight onto each PN axon. **(I)** Correlation of direct excitatory and cascaded inhibitory tuning for LHNs receiving the specified strength of input from either DM1 or DP1m PNs. t-test: ** p < 1×10^-8^. **(J)** Comparison of the correlation between excitatory and inhibitory connectivity and the correlation between predicted excitatory and inhibitory odor tuning for all 970 LHNs in the connectome with at least 5% input from either channel 1 or channel 2 PNs. Colored points denote the cell types examined in panels K and L. Solid red points denote AD1f1, while red circles denote AD1a1. The LH1744-Gal4 line labels both of these types. **(K)** Mean (± s.e.m.) responses to 4 example odors in 4 LHN types before (black) and after (red) application of GABA receptor blockers. Each odor is presented for 2 seconds. Inset: schematic of local application of GABA receptor blockers. N = 16 flies (AD1a1/f1), 11 flies (AV4a1), 11 flies (PD2a1/b1), and 11 flies (PV5a1). Wilcoxon signed-rank test, * p < 0.005 (false discovery rate controlled with Benjamini-Hochberg procedure). **(L)** Mean (±s.e.m.) dF/F responses for each of the 12 tested odors before (abscissa) and after (ordinate) application of GABA receptor blockers, for two LHN types with DM1-dominant input (top) and for two LHN types with DP1m-dominant input (bottom). Green points denote apple cider vinegar and black points denote acetic acid. Sample size is the same as in panel K.

To explore these differences, we first asked how strongly odor responses of LHNs in each channel depend on their full complement of PN inputs. For aversive channel LHNs, the strongest input alone predicted odor coding as well as the weighted sum of all inputs (**Figure 4C**), This is similar to many neurons in the *Drosophila* visual system^36^. Remarkably, appetitive channel LHNs exhibited the opposite pattern: the strongest input alone substantially impaired prediction quality (**Figure 4D**). This inverted trend hinted that the strongest input to some LHNs may be suppressed by inhibition.

Closer examination revealed a striking difference between LHNs with strongest input from DM1 PNs vs. other PNs. Using only the strongest input did not impair prediction quality for all four DM1-dominant LHN types in our sample (although overall predictions were still relatively poor). In contrast, the strongest input model predictions were often negatively correlated with the responses of other LHN types (**Figure 4C, right**). Notably, the biggest mismatches were for LHNs receiving their strongest input from DP1m PNs. We thus suspected that DP1m-dominant LHNs may receive more inhibition than DM1-dominant LHNs.

Local inhibitory LHNs (LHLNs) are major sources of inhibition in the lateral horn^16,37^, which receive direct inputs from PNs and target LHN dendrites and other PN axons. To estimate the inhibition derived from each PN, we computed the cascaded weights of PNs onto LHLNs and LHLNs onto LHNs (i.e., a matrix multiplication between the two layers of weights; **Figure 4E**; Methods). Then, we identified the LHNs that receive substantial direct synaptic inputs (greater than 2.5%) directly from each of the appetitive PNs, and computed the average cascaded inhibitory weight arising from each PN (**Figure 4F**). This revealed DM1 PNs as a major source of cascaded inhibition onto most LHNs, including those with strong excitatory input from DP1m PNs. In contrast, DP1m was not as large of a source of inhibition onto LHNs with strong excitatory input from DM1 (**Figure 4G**). A similar asymmetry was also observed for presynaptic inhibition, where DM1 provides more inhibition onto DP1m PN axons than DP1m provides onto DM1 axons (**Figure 4H**).

This asymmetry in inhibition led to different predicted excitatory and inhibitory odor tuning for LHNs with strong DM1 vs. strong DP1m inputs. Because LHNs with strong direct excitation from DM1 also have strong inhibition originating from DM1, their predicted excitatory and inhibitory input tuning was balanced (i.e., correlated; **Figure 4I**). LHNs with strong excitation from DP1m also have strong inhibition originating from DM1. Because DP1m and DM1 have distinct odor tuning (**Figure 1E**), these LHNs are predicted to receive inhibition that is less balanced with excitation (**Figure 4I**). Inhibition onto DP1m-dominant LHNs is therefore better positioned to alter odor coding than inhibition onto DM1-dominant LHNs.

To test this, we first identified two LHN types predicted to receive inhibition uncorrelated with excitation (AV4a5 and AD1f1), and two LHN types predicted to receive correlated inhibition (PD2a1/b1 and PV5a1; **Figure 4J**). We then measured odor responses before and after applying GABA receptor blockers to the lateral horn (without perturbing the antennal lobe, which is under strong GABAergic regulation^38^; **Figure 4K, S4**). Blockade of inhibition led to odor- and cell-type-specific increases in responses (**Figure 4K**).

Overall, blocking inhibition increased the gain of the two DM1-dominant LHNs (**Figure 4K,L**) but had relatively minimal impact on their odor coding. In contrast, it increased the responses to apple cider vinegar in the two DP1m-dominant LHNs (**Figure 4K,L**), which substantially altered odor coding. Together, these results support the hypothesis of asymmetric inhibition in the lateral horn: DM1 inhibits LHNs driven by other PNs, but other PNs do not inhibit LHNs driven by DM1.

### Unbalanced inhibition contributes to negative valence coding

The DM1 PN strongly encodes positive valence (**Figure 1E**) and also has substantial inhibitory impact in the lateral horn (**Figure 4F,H**). Thus, we reasoned that it may contribute to negative valence coding in LHNs that lack substantial direct excitation from DM1. In support of this idea, the collective odor responses from DP1m-dominant LHNs (which have unbalanced inhibition, **Figure 4J**) correlated negatively with valence, while the DM1-dominant LHNs (which have balanced inhibition, **Figure 4J**) correlated positively (**Figure 5A**). After blocking local inhibition, the correlation between valence and LHN activity vanished for DP1m-dominant LHNs, while it persisted for DM1-dominant LHNs (**Figure 5B**). Accordingly, the net effect of inhibition was positively correlated with valence for DP1m-dominant LHNs, but not for DM1-dominant LHNs (**Figure 5C**). Thus, by selectively suppressing certain odor responses (ACV most prominently), unbalanced inhibition supports negative valence coding.

**Figure 5.**
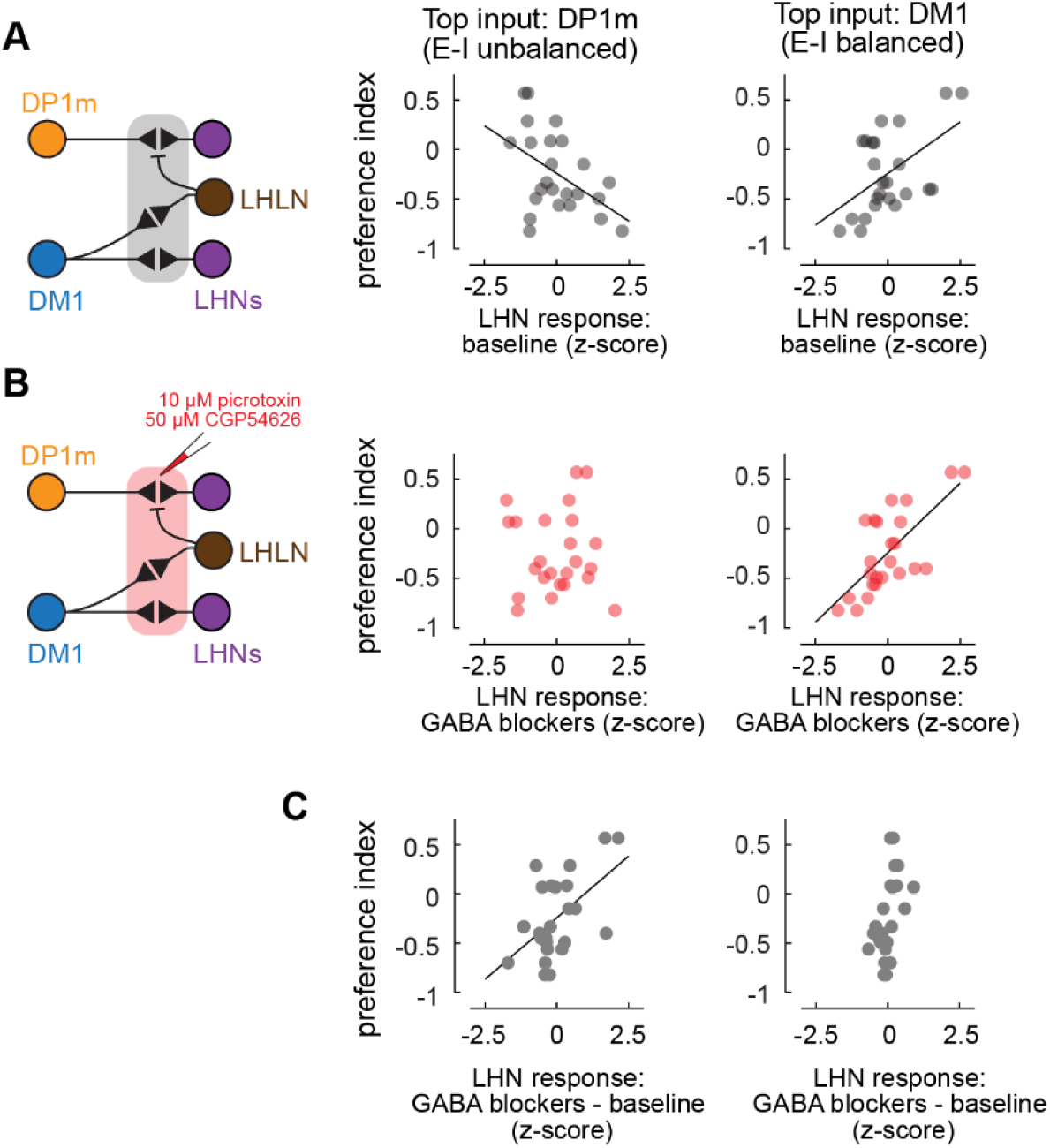
Unbalanced inhibition contributes to negative valence coding. **(A)** Left: schematic of local lateral horn circuitry. Middle: relationship of pooled responses of DP1m-dominant (AV4a5 and AD1a1/f1) LHNs to preference index (Pearson correlation: r = −0.48, p = 0.018). Right: relationship of pooled responses of DM1-dominant (PD2a1/b1 and PV5a1) LHNs to preference index (Pearson correlation: r = 0.52, p = 0.0097). **(B)** Same as (A) but during blockade of inhibition. Middle: Pearson correlation, r = 0.01. p = 0.96. Right: Pearson correlation, r = 0.70, p = 0.0001). **(C)** Net effect of inhibition vs. preference index for DP1m-dominant LHNs (Pearson correlation, r = 0.53, p = 0.0075) and DM1-dominant LHNs (Pearson correlation, r = 0.54, p = 0.0067).

### ACV selectivity arises via integration of acidic and non-acidic odor components

A major contribution to the segregated valence coding in LHNs is the ability of appetitive LHNs to respond more strongly to ACV than to other odors, which is not fully predicted by PN odor coding and PN-LHN connectivity (**Figure 3J**). This mismatch in prediction occurs because these LHNs receive inputs from PNs (e.g., DM1 and DM4) that respond more strongly to odors other than ACV (e.g., ethyl butyrate and ethyl acetate; **Figure 6A**). Although selective local inhibition has been proposed to fine-tune valence coding^14^, our experiments above show that it is not involved, at least in these particular LHNs. How then, do LHNs generate selective ACV responses?

**Figure 6.**
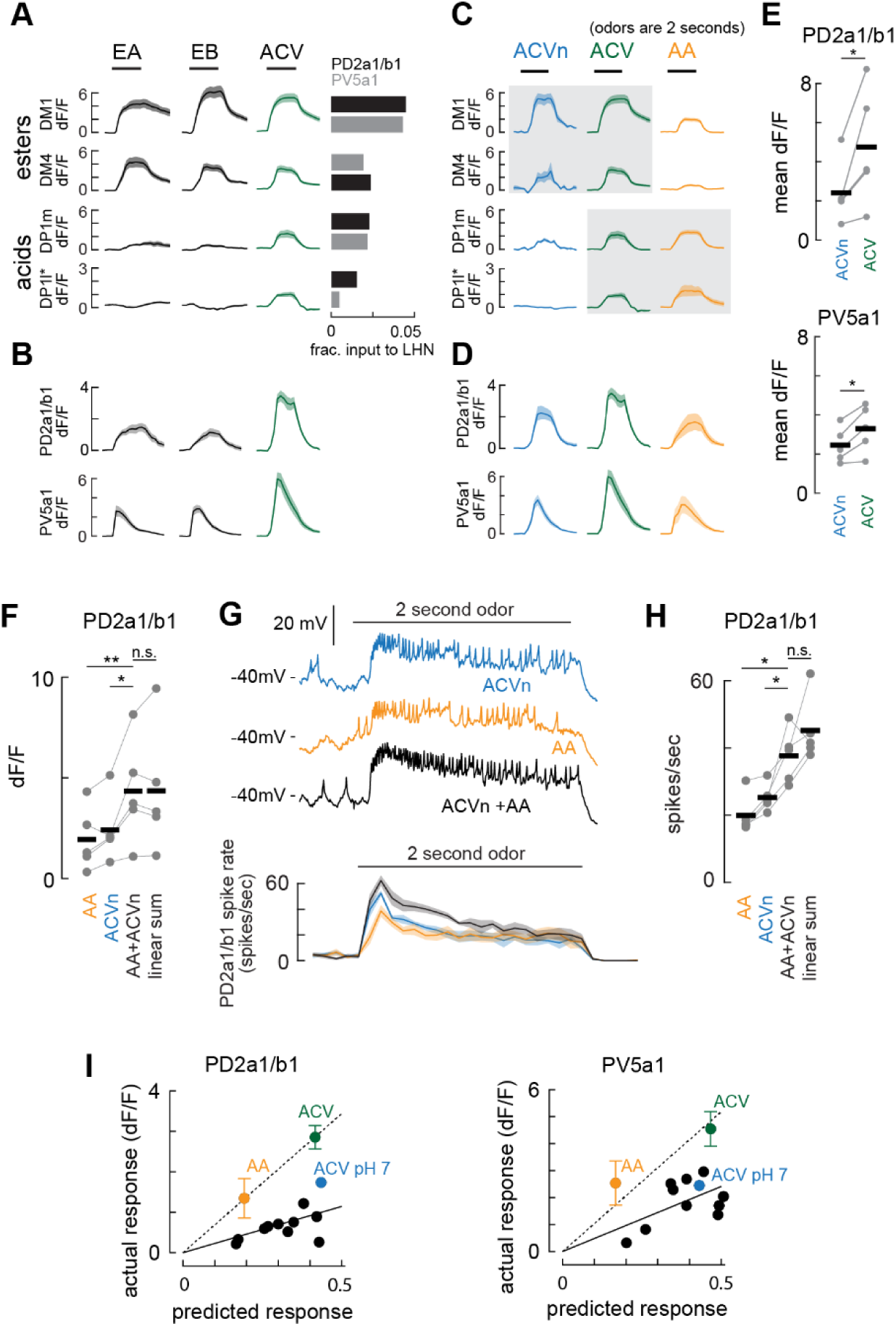
ACV selectivity arises via integration of acidic and non-acidic odor components. **(A)** Mean (± s.e.m.) responses of 2 representative ester-encoding glomeruli and 2 representative acidencoding glomeruli to 2 second presentations of ethyl acetate (EA), ethyl butyrate (EB), and apple cider vinegar (ACV). Right: average fraction of synaptic input (Hemibrain connectome) to PD2a1/b1 and PV5a1 LHNs provided by each of the four PN types. **(B)** Mean (± s.e.m.) fluorescence responses of PD2a1/b1 and PV5a1 LHNs to the same odors as in (A). **(C,D)** Same as (A,B), but for pH-neutralized ACV (ACVn), normal ACV, and acetic acid (AA). ACV responses are repeated from panels A and B for clarity. **(E)** Responses of each fly to ACVn and normal ACV for PD2a1/b1 and PV5a1 LHNs. T-tests, * p < 0.05. **(F)** Responses of each fly to AA, ACVn, AA mixed with ACVn, and the linear sum of the responses to AA and ACVn. Paired t-tests: * p = 0.023, ** p = 0.008. **(G)** Top: example voltage traces from electrophysiological recordings of PD2a1/b1 LHNs to ACVn, AA, and the mixture of the two. Bottom: Mean (± s.e.m.) spike rate during each odor presentation (n = 5 recordings). **(H)** Same as (F), but for average spike rate during the time window between 150msec and 1000msec after valve switching (same recordings as in panel G). Paired t-tests: * p < 0.03. **(I)** Relationship between predicted and actual responses of PD2a1/b1 and PV5a1 LHNs to each odor. Solid lines are linear fits (constrained to pass through the origin) to all odors except AA and ACV. Dashed lines are linear fits (constrained to pass through the origin) to just AA and ACV.

ACV is a complex odor containing both acetic acid and various acetate esters^39^. Notably, acid-responsive PNs – such as DP1m and DP1l – provide synaptic input to ACV-responsive LHNs and respond robustly to ACV (**Figure 1E, 6A**). This raises the possibility that ACV selectivity arises from integration of ester-encoding and acid-encoding PN inputs.

To test this idea, we first neutralized the pH of ACV by adding NaOH (Methods) and measured PN odor tuning. As expected, neutralization preserved responses in DM1 and DM4, but reduced responses in DP1m and DP1l (**Figure 6C**)^40^. In contrast, acetic acid alone activated DP1m and DP1l glomeruli but reduced responses in DM1 and DM4 (relative to ACV). Thus, the acid and ester components of ACV are encoded by separate sets of glomeruli.

We then measured LHN odor responses to these same odors. Neutralized ACV evoked weaker responses than normal ACV (**Figure 6D,E**). This indicates that coactivation of ester-encoding and acid-encoding glomeruli underlies prominent LHN responses. To test this idea further, we compared responses in PD2a1/b1 LHNs to acetic acid, neutralized ACV, and a mixture of these two odors. The mixture elicited responses that were indistinguishable from the linear sum of the responses to each odor alone (**Figure 6F**). This was neither due to intrinsic nonlinearities of GCaMP nor possible heterogeneities of LHNs within these types, because we observed a nearly identical and highly consistent effects in individual LHN spike rates (**Figure 6G,H**).

The simple linear feedforward model predicted that ACV and acetic acid activate LHNs with a higher gain than other odors (**Figure 6I**). This may occur via stronger synapses from acid sensing glomeruli, possibly involving different forms of short-term plasticity or different postsynaptic receptor types. Alternatively, ester-responsive glomeruli may be subject to greater normalization in the lateral horn, which suppresses their impact on LHNs when driven broadly.

### LHN adaptation is orthogonal to valence

Sensory codes can also utilize time as a dimension by encoding aspects of a sensory stimulus in the relative timing or adaptation profiles across a population of neurons. Some LHNs are known to adapt to ongoing odor stimulation in different ways. PD2a1/b1 LHNs adapt modestly, faithfully encoding the continuing presence of odor; PV5a1 LHNs adapt strongly, encoding only the initial exposure to odor^15^. Because these two cell types have similar odor tuning, it raises the possibility that LHN adaptation varies independently of valence coding across the population. To explore this idea, we directly investigated odor response dynamics in each LHN type in our dataset.

We first confirmed the distinct adaptation of PD2a1/b1 and PV5a1 LHNs. PD2a1/b1 adapted modestly, with calcium levels relaxing to a level elevated above baseline for the duration of a 10 second odor pulse (**Figure 7A**). In contrast, PV5a1 adapted completely, with calcium returning to baseline levels within the first few seconds of a 10 second odor. Although calcium does not fully settle to steady state levels within our typical 2 second odor stimulation, clear differences in adaptation are still evident (**Figure 7B**). Thus, differences in calcium dynamics for 2 second odor stimulation reflect differences in adaptation between LHN types.

**Figure 7.**
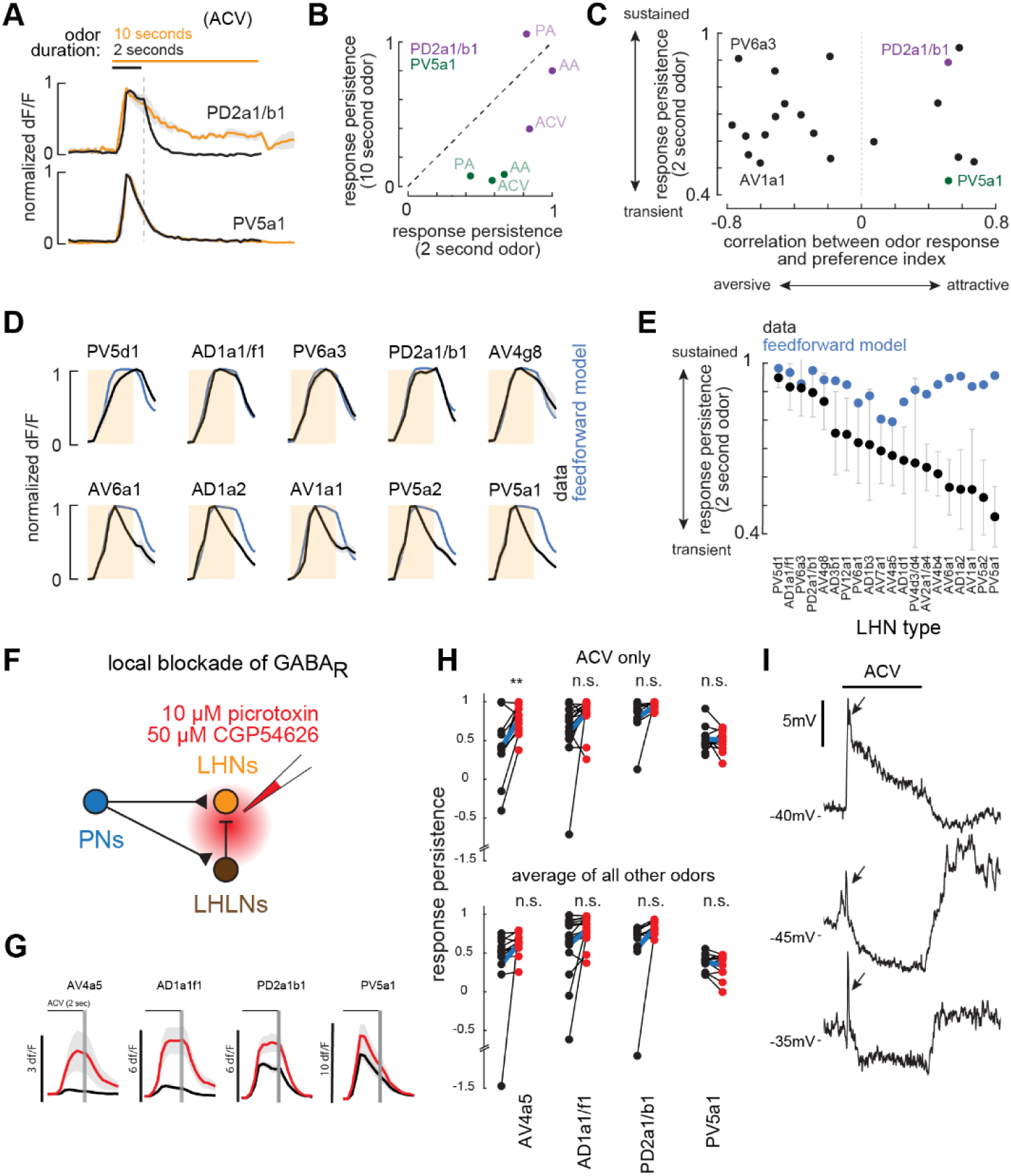
LHN adaptation dynamics form an orthogonal coding dimension to valence. **(A)** Mean (± s.e.m.) responses to 2 seconds (black) and 10 seconds (orange) of ACV in PV5a1 (n=4) and PD2a1/b2 (n = 5). Responses do not reach steady state after only 2 seconds, but the distinction in adaptation is already clear. **(B)** Comparison of response persistence after 2 seconds and after 10 seconds, measured separately for three odors for PV5a1 and PD2a1/b1 LHNs. AA: acetic acid, PA: pentyl acetate, ACV: apple cider vinegar. **(C)** The strength of LHN valence coding (correlation between neural response and behavioral response) compared to 2-second response persistence for each LHN type. There is no significant relationship between valence coding and response persistence. Shuffle test (1000 shuffles), p = 0.5560. **(D)** LHN response dynamics (mean ± s.e.m. across all odors normalized to their peak responses) compared with average response dynamics predicted from the linear feedforward model, for 10 LHN types. **(E)** Comparison of actual vs. predicted response persistence in model vs. actual LHN responses. Error bars are standard deviation across odors. **(F)** Schematic of local GABA antagonism in the LH to assess LHN dynamics. **(G)** Blockade of local inhibition affects ACV response dynamics in AV4a5 LHNs, but not for other odors or for other LHN types. **(H)** Top: comparison of response persistence for ACV responses before (black) and after (red) application of GABA receptor blockers (paired t-tests: AV4a5, p= 0.0102; AD1a1/f1, p= 0.1083; PD2a1/b1, p= 0.1169; PV5a1, p=0.6271). For all other odors, dynamics did not change with GABA antagonism (paired t-tests: AV4a5, p= 0.2757; AD1a1/f1, p= 0.2202; PD2a1/b1, p =0.2757; PV5a1, p = 0.2757). **(I)** Voltage responses (each an average of 5 traces) from electrophysiological recordings of three AV4a5 LHNs in response to 2 second stimulation with apple cider vinegar.

Across the 20 LHN types we sampled, we observed a broad range of dynamics, including both sustained and transient types which encoded aversive odors (**Figure 7C**). We observed no correspondence between LHN adaptation profile, and valence coding. Thus, LHN adaptation is a coding dimension that is orthogonal to valence.

In principle, LHNs could inherit their adaptation dynamics from their PN inputs, or they could be computed *de novo* in the lateral horn. While some PNs do adapt relatively strongly, we observed mostly sustained responses from the PNs making strong connections to the LHNs in our sample (**Figure S7**). We thus used the feedforward linear model to predict the dynamics of each LHN’s odor responses from the dynamics of its complement of PN inputs. This model recapitulated the dynamics of sustained LHNs, but not the transient LHNs (**Figure 7D,E**). Together, these findings suggest that sustained LHN responses are inherited from PNs, while transient LHN responses are computed *de novo*.

Adaptation in some LHNs is known to arise from target-cell specific short term plasticity of PN presynapses^15^. However, local inhibition may also contribute. We thus asked how local inhibition (**Figure 7F**) impacts LHN odor response dynamics. In most cases, blockade of inhibition did not substantially alter odor dynamics (**Figure 7G,H**). The singular exception was the response of AV4a5 to ACV (**Figure 7H**). Inhibition strongly suppresses this response, leaving only a small calcium transient (**Figure 7G**). To investigate the underlying membrane dynamics of this transient response, we made patch-clamp recordings from AV4a5 LHNs. These revealed that ACV drove very brief initial depolarizations that were sometimes followed by sustained hyperpolarization (**Figure 7I**). This is compatible with the additional synaptic delay of inhibition relative to excitation. Together, this demonstrates that local inhibition can generate transient LHN dynamics but is a more odor-specific mechanism than short-term plasticity.

## DISCUSSION

### Segregated pathways for valence

Excitatory feedforward convergence in sensory processing must strike a delicate balance: it is needed for assembling receptive fields, but it also irreversibly mixes information. Excitatory connectivity is therefore tightly regulated so that convergent neurons usually carry functionally related information^41,42^. The first stage of the olfactory system is an extreme example of this, where functionally identical ORNs exclusively converge onto the same PNs^43–46^, maintaining the integrity of each olfactory channel. The connections that *don’t* exist are just as important as those that do, by preventing haphazard interference between information channels.

Our results show that olfactory valence coding in *Drosophila* follows this principle. PNs encoding different odors of the same valence preferentially converge onto the same LHNs, sacrificing information about exact odor identity in the service of building a more explicit neural code for instinctive olfactory judgements. Similar to the antennal lobe, this convergence closely tracks anatomy: PNs encoding attractive odors project to a distinct lateral horn subregion from PNs encoding aversive odors and pheromones^12,34,35^. Thus, odor valence maps onto spatial subregions of the lateral horn, despite the lack of insulated compartments defined by ensheathing glia^47^. These results are compatible with prior evidence of valence coding^14,48^ and evidence of spatial segregation of odor representations^49^ in the lateral horn.

Feeding, mating, and avoiding danger rely heavily on innate odor valence, but also benefit from prior experience. Learned odor associations are formed in the mushroom body, where PNs converge onto KCs. Compared to PN-LHN convergence, PN-KC convergence is less structured^13,31–33^, which contributes to a higher dimensional odor representation^50,51^. We find that the modest structure that does exist in PN-KC connectivity has the opposite relationship to innate valence as PN-LHN connectivity: convergence tends to separate odor representations with similar valence. This complementary architecture should support the ability to learn to discriminate between subtle differences of PN activity patterns^29^.

### Connectivity and computation underlying valence coding

Our results revealed a curious feature of the DM1 PN: despite having the strongest positive valence association of any PN, it also has the widest-ranging local inhibitory impact of any PN. Because DM1 ORNs make the largest causal contribution to odor attraction of any individual ORN type^17,24,52^, it is at first puzzling that their cognate PN is the source of so much inhibition in the lateral horn. However, DM1’s strongest (polysynaptic) inhibitory connections target different LHNs than its strongest (monosynaptic) excitatory connections. Thus, its anatomical projective field forms a “center-surround” arrangement onto LHNs, similar to circuitry in the vertebrate retina^53,54^.

LHNs in the center of DM1’s projective field generally have balanced excitatory and inhibitory inputs, because DM1 is a major supplier of both. Inhibition scales the gain of these LHNs, but does not significantly change their odor and valence tuning, consistent with classical models of balanced inhibition^55^. In contrast, LHNs in the surround of DM1’s projective field generally have unbalanced excitatory and inhibitory inputs, because DM1 mostly provides inhibition. Inhibition significantly changes the odor tuning of these LHNs, consistent with previous reports of odor specific inhibition in select LHN types^37^. Because DM1 encodes positive valence, its inhibitory impact contributes to negative valence coding.

This configuration provides two complementary advantages for valence coding. First, the projective field surround limits the impact of other PNs that encode valence less directly. For example, AV4a5 receives strong input from DP1m PNs, which respond to ACV, but also from some PNs encoding aversive odors. Inhibition specifically suppresses responses to ACV, resulting in a more consistent negative valence code. Second, the projective field center creates the opportunity to sharpen selective responses to attractive odors. Because DM1 inhibition is offset by excitation in these LHN types, they are more free to integrate DM1 activity with that of other glomeruli, strengthening responses to ACV, and contributing to overall encoding of positive valence.

We found that both PD2a1/b1 and PV5a1 both linearly integrated acid and ester components of ACV, leading to stronger responses than for other odors driving similar levels of feedforward input. Thus, the projective field center amplifies responses to ACV while the projective field surround attenuates them. The exact mechanism establishing the selectivity of amplification, however, remains unknown.

Our results stand in contrast to a recent study which concluded that balanced inhibition in LHNs is necessary for positive valence coding, by selectively suppressing responses to specific odors (such as ethyl butyrate, which drives strong PN activity, but does not have a strong innate valence)^14^. Because that study recorded from LHN somata, they were unable to link measured odor responses to connectivity (because soma location does not definitively identify LHN type). Their conclusions were instead based largely on a computational model of inhibition in the lateral horn. By experimentally manipulating inhibitory signaling while measuring odor responses from genetically identified LHN types, our results instead suggest that balanced inhibition has little impact on valence coding. However, our sample size is restricted by available Gal4 lines, so inhibition may have a larger impact on other LHN types which we did not study here.

### Valence coding and adaptation

Adaptation enables sensory neurons to continuously adjust their sensitivity to match current stimulus statistics^56^. This can make coding more efficient but also introduces ambiguities in the mapping between stimulus and response that can be difficult to resolve. Our data suggest that the olfactory system may limit ambiguities via parallel processing, by representing the same odor information in different neurons with different adaptation properties, a configuration reminiscent of retinal circuits^57^.

In most cases, blocking inhibition did not affect strongly adapting odor responses. This suggests a widespread role for target-neuron specific short-term plasticity in implementing diverse adaptation throughout the lateral horn, a phenomenon we previously demonstrated at two PN-LHN synapse types^15^. In one case (ACV responses in AV4a5 LHNs) blocking inhibition transformed a transient response into a more sustained response. Inhibition-mediated transient responses can be extremely brief due to the short time delay of polysynaptic inhibition^58^. Thus, there are at least two distinct mechanisms for implementing response transience in the lateral horn.

Splitting olfactory pathways by both valence and dynamics likely supports innate behavioral responses. Flies rapidly respond to dynamic odors but also maintain responses to sustained odor^15,59^. Separate LHNs encoding relative and absolute odor properties should enable downstream circuits to regulate different motor programs to make appropriate behavioral responses. This arrangement could also enable pathway-specific processing, for instance by selectively amplifying transient representations to make quick turns, or by integrating odor information over time to make more accurate judgements (similar to what occurs in the mushroom body^60^). Investigating how these dynamics integrate into downstream circuits, such as the fan shaped body and superior protocerebrum^22,61^, should provide an opportunity to test the functional role of pathway splitting for both neural computation and behavior.

### Limitations of this study

While we measured odor coding from a nearly complete subset of PNs, an important limitation of this study is the limited sample size of LHN types. The range of valence coding we observed was statistically indistinguishable from a larger sample measured from LHN somata^14^, but it remains possible that unobserved LHN types operate differently from those in our sample. Because neuron identity is crucial for mapping structure to function, generating more comprehensive collections of driver lines to target more LHN types is an important goal for the future. Our odor panel spanned the range of aversive and appetitive valence, but every physiology experiment is limited by the number of stimuli that can be presented. Given the special integration that we observed for ACV, (a natural mixture of many individual volatile chemicals), it will be useful to explore how LHNs encode a broader range of odor mixtures, especially those that combine esters and acids in natural ratios. Finally, complex odor dynamics, like those that occur in natural odor plumes, may engage LHN adaptation differently from our static odor pulses. Future work exploring the temporal dynamics of LHNs to these kinds of stimuli may reveal more specific pathways with heterogeneous adaptation properties.

## Acknowledgements

We thank Fernando J Figueroa Santiago for help with earlier connectomic analyses on inhibitory circuits in the lateral horn, Poonam Mishra for assistance with picospritzer set up, and members of the Jeanne Lab for helpful discussions and for comments on the manuscript. We also thank Joel Greenwood and Paul Shamble of the Yale Neurotechnology Core for their technical expertise and assistance. This work was supported by NIH grants R01 DC018570 and R01 NS116584, the Richard and Susan Smith Family Award for Excellence in Biomedical Research, the Klingenstein-Simons Fellowship Award in Neuroscience, and an innovative research award from the Kavli Institute for Neuroscience at Yale University to J.M.J. K.M.L. was supported by a James Hudson Brown-Alexander Brown Coxe Postdoctoral Fellowship in the Medical Sciences at Yale University School of Medicine and NIH fellowship F32 DC019521.

## KEY RESOURCES TABLE

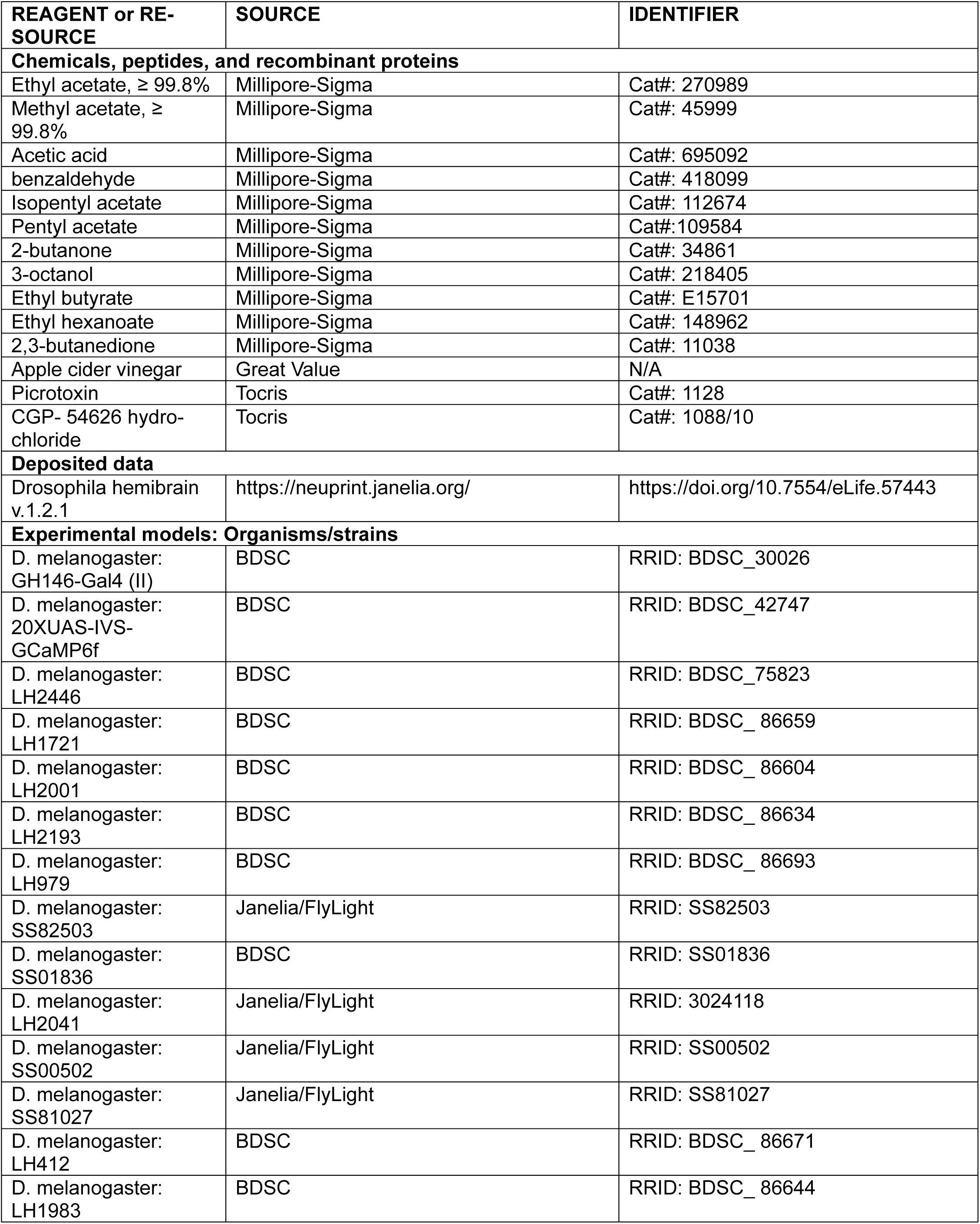
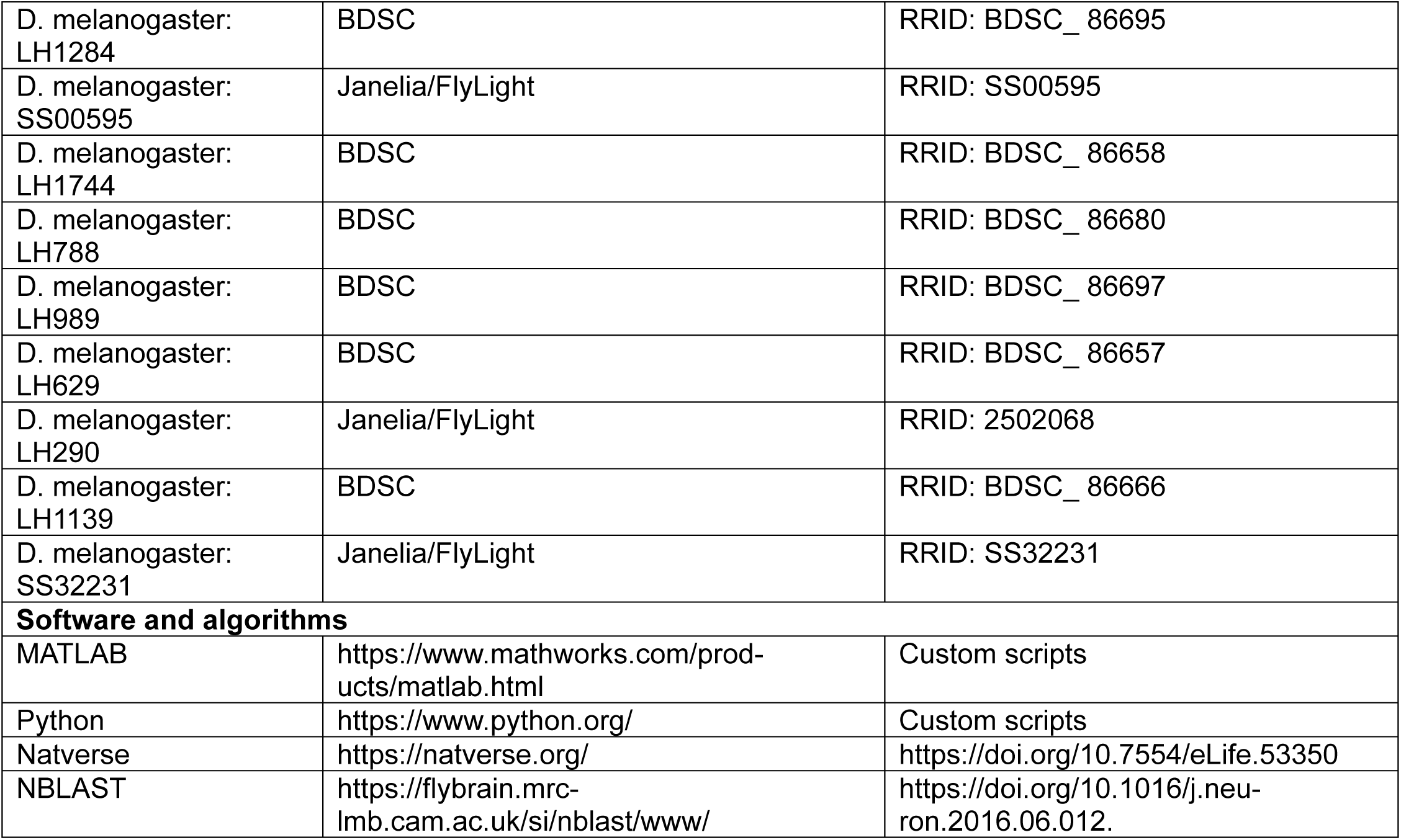

## RESOURCE AVAILABILITY

### Lead Contact

Further information and requests for resources and reagents should be directed to and will be fulfilled by the lead contact, James M. Jeanne.

### Materials Availability

This study did not generate new unique reagents.

### Data and code availability

Imaging data will be deposited and publicly available as of the date of publication. Accession numbers are listed in the key resources table. All original code will be deposited at Zenodo and is publicly available as of the date of publication. DOIs are listed in the key resources table. Any additional information required to reanalyze the data reported in this paper is available from the lead contact upon request.

## EXPERIMENTAL MODEL

Flies (*Drosophila melanogaster*) were raised on conventional cornmeal agar medium or German food formulation under a 12 h light, 12 h dark cycle at 25°C. Calcium imaging experiments were performed on fed adult female flies 1-4 days after eclosion. All flies used for behavior experiments were in the IsoD1 genetic background^62^. Behavior experiments were performed on fed adult male and female flies 3-4 days after eclosion.

## METHOD DETAILS

### Fly preparation for imaging and electrophysiology

For imaging and electrophysiology experiments, each fly was cold-anesthetized, positioned into a small holder made of stainless steel shim stock (0.001″ thick), and affixed into position using paraffin wax such that the antennae were under the foil and dry while the head and brain was bathed in external saline. The external saline contained (in mM): 103 NaCl, 3 KCl, 5 N-tris(hydroxymethyl) methyl-2-aminoethane-sulfonic acid, 8 trehalose, 10 glucose, 26 NaHCO_3_, 1 NaH_2_PO_4_, 1.5 CaCl_2_ and 4 MgCl_2_ (osmolarity adjusted to 270–275 mOsm). A small window in the top of the head capsule was dissected using electrolytically sharpened tungsten wires and fine forceps. Fat, air sacs, and trachea were removed from above the brain.

For electrophysiology experiments, fine forceps were then used to gently remove the perineurial sheath only above the area of the brain housing the target somata. Large-bore cleaning pipettes were used to remove residual glia and interfering somata to gain clear access to target somata. External saline was bubbled with 95% O_2_ and 5% CO_2_ and reached an equilibrium pH of 7.3.

### Calcium Imaging

*In vivo* calcium imaging of PNs and LHNs was performed on a 2-photon laser scanning microscope (Scientifica MP-2000) using a 20x 1.0NA water-immersion objective lens (Olympus). Either a Titanium-Sapphire laser (Coherent Chameleon Ultra I) tuned to 920nm or a fixed wavelength 920nm laser (Spark Alcor XSight) was used to excite GCaMP6f, and fluorescence emission was collected with a GaASP PMT, controlled by ScanImage software (MBF Bioscience). Images were collected at an acquisition rate of 4.22 frames per second. For PN imaging, measurements were collected sequentially from multiple planes spanning the antennal lobe for each fly. Glomeruli were identified post-hoc by comparing baseline fluorescence patterns with established atlases ^9,63^. For LHN imaging, measurements were collected from a single plane containing the largest branching of neurites in the lateral horn. dF/F was calculated as the increase in fluorescence per glomerulus or LHN during odor presentation, normalized by the baseline fluorescence in the corresponding glomerulus or LHN.

### Electrophysiology

Recordings were obtained using an Olympus BX51 upright microscope with 40X water immersion objective. One LHN was recorded per fly. Patch-clamp electrodes were filled with an internal solution of (in mM): KCH_3_SO_3_H 160, HEPES 10, MgATP 4, Na_3_GTP 0.5, EGTA 1, biocytin hydrazide 13 (pH = 7.3, adjusted to 265mOsm). Patch pipettes were made from borosilicate glass (Sutter; 1.5-mm outer diameter, 0.86-mm inner diameter) and were pulled and pressure polished to create a relatively long taper with final pipette tip opening of about 0.75μm in diameter ^64^. Recordings were obtained with an Axopatch 200B model amplifier in current clamp mode with CV-203BU head stages, digitized at 10-Hz with an analog-to-digital converter (National Instruments), and saved to disk using the MATLAB data acquisition toolbox. Recordings were not corrected for a liquid junction potential of ∼13mV ^65^.

Following most LHN recordings, brains were processed for immunohistochemistry exactly as described previously ^13^, and imaged with a Zeiss LSM 880 confocal microscope to visualize the biocytin fill and confirm the LHN type identity.

### Odor preparation and delivery

For all imaging and electrophysiology experiments, a clean air stream (1360 mL/min) was filtered through activated carbon and directed to the fly through a carrier tube. Separate air streams of 12 ml/min were directed under the control of solenoid valves (The Lee Company, model LHDA1231415H) into the headspace of 2mL vials (Thermo Scientific, National C4011-5W) containing odors. The odor streams joined the carrier stream 7-8.5cm from the end of the tube. All imaging and electrophysiology experiments used the odor configuration in Figure S2B of Kim et al.^15^. Here, PTFE tubing of 9 cm and 13 cm lengths were used to connect the odor vial to the solenoid valve, and to the carrier stream respectively. Odors were diluted to 1% concentration in water, except ACV and acetic acid, which were undiluted. Identical odor concentrations were used for behavior experiments. To ensure sufficient headspace in each odor vial, the final volume of odor dilutions was 0.5 mL. The timing of valve opening and closing was controlled by a custom MATLAB script. Each odor pulse was controlled by opening the valve for 2sec, except for the experiments in Figure 7A, where valves were opened for 10 sec.

For neutralizing ACV, a stock solution of NaOH was prepared by adding solid NaOH pellets to water. Concentrated NaOH was titrated into undiluted ACV until pH reached neutral 7. The volume of NaOH solution added to neutralize ACV was recorded so that an exact volume of water could be added to a separate undiluted ACV vial, to ensure reduced responses in PNs were not purely due to dilution by the added water in the NaOH solution.

### Pharmacology and Pressure injection of GABA antagonists

CGP54626 hydrochloride (Tocris) and picrotoxin (Tocris) were prepared as concentrated stock solutions dissolved in DMSO, then diluted in physiological saline to a final concentration of 50μM CGP, and 10μm picrotoxin. The solution of antagonists was loaded into a pulled glass pipette made of borosilicate glass with a tip diameter of 2-3uM and positioned onto either the antennal lobe or the lateral horn neuropil. After recording baseline responses to all 12 odors, four 400ms pulses of drugs were pressure ejected (Picospritzer III, Parker Hannifin) using 10 psi. After waiting 2 minutes for drugs to take effect, responses to all 12 odors were recorded again. This protocol was sufficiently local as to not impact odor processing in the antennal lobe (**Figure S4**).

PN responses were similarly affected by local and brain-wide (bath) application of antagonists at identical concentrations, indicating that our disruption of GABA signaling in the lateral horn is equivalent to bath application at the same concentration. Following GABA antagonism, dF/F for each LHN was calculated as the increase in fluorescence during odor presentation, normalized by the pre-drug baseline fluorescence, with the change in baseline due to antagonism subtracted. This was done to assess how antagonism changes odor responses independent of tonic effects on the baseline fluorescence.

### Behavioral place preference assay

Behavioral odor preferences were tested in a custom aluminum rectangular arena (115 x 30mm) maintained at 32°C, with odor delivered from inlets on either end. A constant airstream of 420 mL/min was maintained from each port, with odor pulses generated by using computer-controlled solenoid valves to divert 120 mL/min of the airflow through an odor vial to replace an equal fraction of clean air, keeping the total flow from that port constant. A vacuum (1200mL/min) in the center of the arena cleared out odors between trials. Odor vials consisted of 50mL Falcon tubes fitted with luer connectors, with concentrations matched to those used in imaging experiments. Groups of 8–12 fed flies (3-4 days old) were introduced without anesthesia and allowed to acclimate for 2 minutes before trials. Each trial consisted of a 30 second baseline, a 90 second odor stimulus, and a 30 second post-stimulus period. Each cohort of flies experienced separate trials with the odor presented from the top and bottom inlets, with equal numbers of each type. Behavior was recorded from above at 3 fps under IR LED illumination, and fly positions were extracted using SLEAP, a deep learning-based multi-animal pose tracking framework^66^ trained in-house on manually annotated data. A preference index was calculated at each time point as (*N_odor_* – *N_air_*)/*N_total_*, where *N_odor_* and *N_air_* are the numbers of flies in the halves of the arena closer to the odor and air inlets, respectively. For each cohort, the mean of the last 30 seconds of odor presentation was computed for the top and bottom trials and averaged together. The reported odor preference index was then calculated as the mean of the cohort preference indices across all cohorts tested.

## QUANTIFICATION AND STATISTICAL ANALYSIS

### Connectome analysis

Synaptic connectivity data were obtained from the Hemibrain connectome (version 1.2.1) using custom Python scripts and the neuprint-python API. Synapse counts between all canonical cholinergic uniglomerular PNs (those in the adPN or lPN lineages) and every LHN were retrieved with “fetch_adjacencies.” Raw synapse counts per connection were normalized by the total number of input synapses on each LHN instance.

Local neurons in the lateral horn are all predicted to be either GABAergic or glutamatergic^16,67^, both of which are expected to be inhibitory in the fly olfactory system^68^. Some local neurons have processes which modestly spill out from the anatomical lateral horn boundary of the Hemibrain. To accommodate these, we assumed that all LHNs with more than 60% of their output synapses in the lateral horn neuropil were local neurons, and thus inhibitory. To characterize each PNs inhibitory impact on each LHN, we summed the product of PN-LHLN and LHLN-LHN connection weights over all LHLNs.

For comparison of connectome predicted LHN odor responses vs. actual LHN odor responses, we pooled over all LHNs in the connectome of the type expressed by each Gal4 line we used for imaging. Note that LH788-Gal4 labels AV4a5 LHNs, which is different from its published annotation (**Figure S1**).

### Imaging and electrophysiology data analysis

Average PN responses to each odor were computed over a 1.2 second window starting 1 second after solenoid valve switching. The delay corresponds to the time for the odor molecules to propagate from the odor vial to the fly. Average LHN responses to each odor were computed over a 1.25 second window starting 0.75 seconds after solenoid valve switching. Adaptation (response persistence) was quantified as the ratio of the dF/F value at the end of odor delivery to the dF/F value at the onset of odor delivery. Spikes in electrophysiological recordings from PD2a1/b1 LHNs were detected exactly as described previously^15^. Average spike rates were computed during the first 1 second of the odor response.

### PN and LHN activity vector analysis

Average PN and LHN activity vectors were computed by assembling all PN dF/F responses to each odor into a single vector. The Euclidean distance between pairs of vectors is then the length of the line connecting them in 41-dimensional space (PNs) or 20-dimensional space (LHNs). Activity vectors for the linear projection of PN activity into simulated LHN space were computed by matrix multiplication of the PN activity vectors for each odor with the matrix of PN-LHN connection weights. Because there are 1476 identified LHNs in the Hemibrain connectome dataset, this yielded a 1476-dimensional space. Random projections of simulated LHN space were obtained by shuffling the entries in the PN-LHN connectivity matrix. An identical procedure was followed to analyze simulated KC activity vectors, which occupied 1828-dimensional space.

### PN convergence analysis and channel identification

Each entry in the PN-PN correlation matrix in Figure 3A was computed as the Pearson correlation coeffencent between the connectivity vectors for each pair of PNs. We only considered the 1370 LHNs in the Hemi-brain which had positive connectivity with one of the 48 PN types of either at anterodorsal or lateral lineage.

Hierarchical clustering was performed using the MATLAB functions “pdist” (with a cosine distance metric) and “linkage” (with distance between clusters computed as the unweighted average distance). A cosine linkage distance of 0.946 was used to separate PNs into distinct channels. LHNs with significantly biased inputs from channel 1 or channel 2 were identified with a chi-square test compared to a Bernoulli distribution with p = 0.5. The false discovery rate was controlled via the Benjamini-Hochberg procedure. Total projected activity onto channel 1 and channel 2 LHNs, and Euclidean distances were computed as above.

### Predicted inhibition-excitation odor tuning correlation

Predicted excitatory inputs were simulated as the connectome weighted sum of the odor responses from all PN inputs to each LHN. Predicted inhibitory inputs were simulated in an identical manner, except we used the cascaded matrix computed from the PN-LHLN and LHLN-LHN connection weights. The odor tuning correlation for each LHN was then evaluated as the Pearson correlation coefficient of these two sets of predicted odor responses.

