## Supplemental information for "Sensory processing reformats odor coding around valence and dynamics"

**Table S1**

| Fly line | genotype | Cell type labeled |
|---|---|---|
| GH146 | y[1] w[1118]; P{w[+mW.hs]=GawB}GH146 | PNs |
| VT033006 | w[1118]; P{y[+t7.7] w[+mC]=VT033006-GAL4.DBD}attP2 | PNs |
| IR75a | w[*]; P{w[+mC]=Ir75a-GAL4.S}BT12.1/TM6B, Tb[1] | DP1l ORNs |
| LH2446 | w; VT029753-p65ADZp in attP40; 55C09-ZpGdbd in attP2 | AD1d1 |
| LH1721 | w; 92D09-p65ADZp in attP40/CyO::Tb-RFP; 85F05-ZpGdbd in attP2/TM6B | AD1a2 |
| LH2001 | w; VT022244-p65ADZp in attP40/CyO::Tb-RFP; VT014480-ZpGdbd in attP2/TM6B | AV7a1 |
| LH2193 | w; VT060077-p65ADZp in attP40/CyO::Tb-RFP; VT029317-ZpGdbd in attP2/TM6B | PV6a3 |
| LH979 | w; 84F07-p65ADZp in attP40; 37A11-ZpGdbd in attP2 | PV4d3/d4 |
| SS82503 | w; R64A11-p65ADZp in JK22C/CyO,Tb; R65G11-ZpGdbd in attP2 | AD1b3 |
| SS01836 | w; VT014093-p65ADZp in attP40/CyO,Tb; 95A10-ZpGdbd in attP2 | AV4g8 |
| LH2041 | w; VT049488-p65ADZp in attP40/CyO::Tb-RFP; VT043152-ZpGdbd in attP2/TM6B | PV12a1 |
| SS00502 | w; 25B07-p65ADZp in attP40; 45F12-ZpGdbd in attP2 | PV5a2 |
| SS81027 | w; 78B06-p65ADZp in attP40; VT037023-ZpGDBD in attP2 | PV6a1 |
| LH412 | w; 22B12-p65ADZp in attP40; 54G12-ZpGdbd in attP2 | AV4b4 |
| LH1983 | w; 76E07-p65ADZp in attP40/CyO::Tb-RFP; VT008671-ZpGdbd in attP2/TM6B | AV1a1 |
| LH1284 | w; 38F08-p65ADZp in attP40; 52H12-ZpGdbd in attP2 | AD3b1 |
| SS00595 | w; 52H01-p65ADZp in attP40; 12G04-ZpGdbd in attP2 | PV5d1_a |
| LH1744 | w; 92D09-p65ADZp in attP40/CyO::Tb-RFP; 92A07-ZpGdbd in attP2/TM6B | AD1a1/f1 |
| LH788 | w; 34C08-p65ADZp in attP40; 22C12-ZpGdbd in attP2 | AV4a5 |
| LH989 | w; 29G05-p65ADZp in attP40; 37G11-ZpGdbd in attP2 | PD2a1/b1 |
| LH629 | w; 30A10-p65ADZp in attP40/CyO::Tb-RFP; 10H02-ZpGdbd in attP2/TM6B | AV2a1/a4 |
| LH290 | w; 38D01-p65ADZp in attP40/CyO; 45F12-ZpGdbd in attP2/TM2 or TM6B | PV5a1 |
| LH1139 | w; 93A02-p65ADZp in attP40/CyO::Tb-RFP; 44G08-ZpGdbd in attP2/TM6B | AV6a1 |
| SS32231 | w; VT040847-p65ADZp in attP40; VT000772-ZpGdbd in attP2 | AV2c1 |

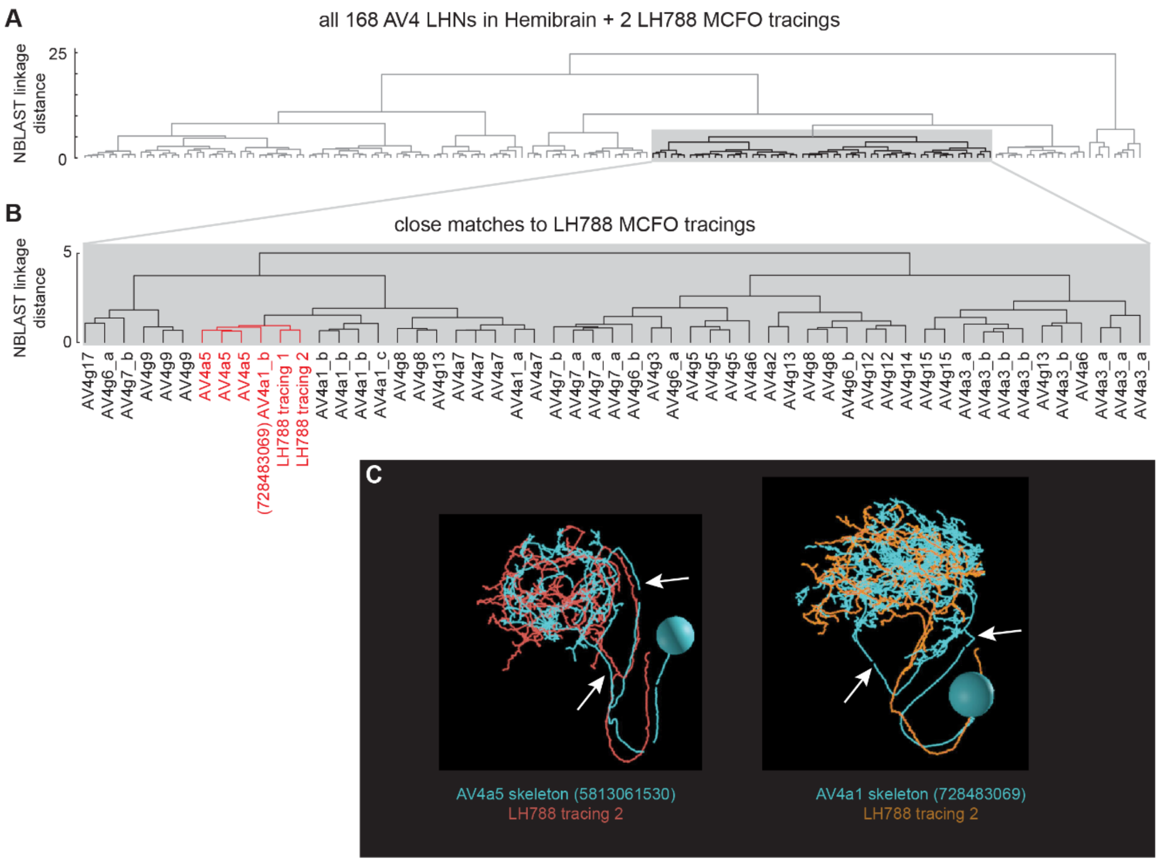


**Figure S1. LH788-Gal4 Labels AV4a5 LHNs**

**(A)** NBLAST (Costa et al., 2016) analysis between all AV4 LHNs (168 total) identified in the hemibrain, and tracings of two Multicolor Flip-out clones from LH788-Gal4 (Nern et al., 2015). Shaded region identifies closest matches to MCFO tracings.

**(B)** NBLAST identifies AV4a5 and AV4a1_b as closest matches to MCFO tracings.

**(C)** Detailed comparison of the neurite projections of AV4a5 and AV4a1_b. The MCFO tracing and AV4a1_b exhibit a mismatch in primary neurite tract, while neurite tract of LH788 tracing and AV4a5 match closely.

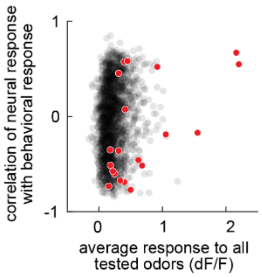


**Figure S2. The odor coding valence of the 20 LHN types studied here is representative of larger populations.**

Comparison between total response strength and the odor coding valence (correlation between neural response and behavioral response across odors). Red points are the 20 LHN types (based on specific Gal4 drivers) in this study. Black points are the 2062 individual LHN somata from Someya et al. (2025). Because our study imaged from LHN dendrites instead of somata, the responses tend to be larger.

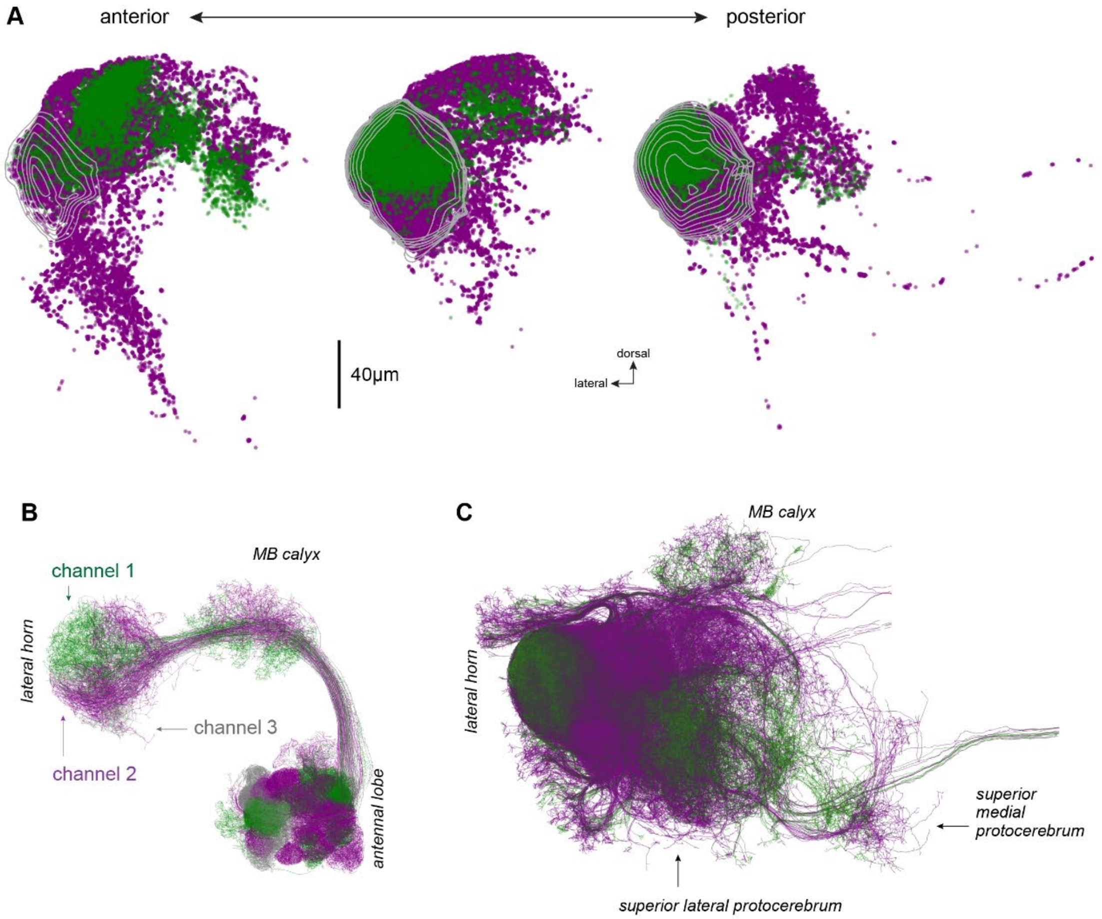


**Figure S3. Channel 1 and 2 PNs and LHNs have distinct anatomy.**

**(A)** Comparison of channel 1 (green) and channel 2 (purple) LHN synapse locations. Channel 1 (appetitive) LHNs target anterodorsal regions of the superior protocerebrum. Channel 2 (aversive) LHNs target posterodorsal regions of the superior protocerebrum and also the ventrolateral protocerebrum. Contours denote the lateral horn neuropil, as in Figure 3B.

**(B)** Oblique view of skeletons of PNs in channel 1 (green), channel 2 (purple), and channel 3 (gray).

**(C)** Same as (B) but for LHNs.

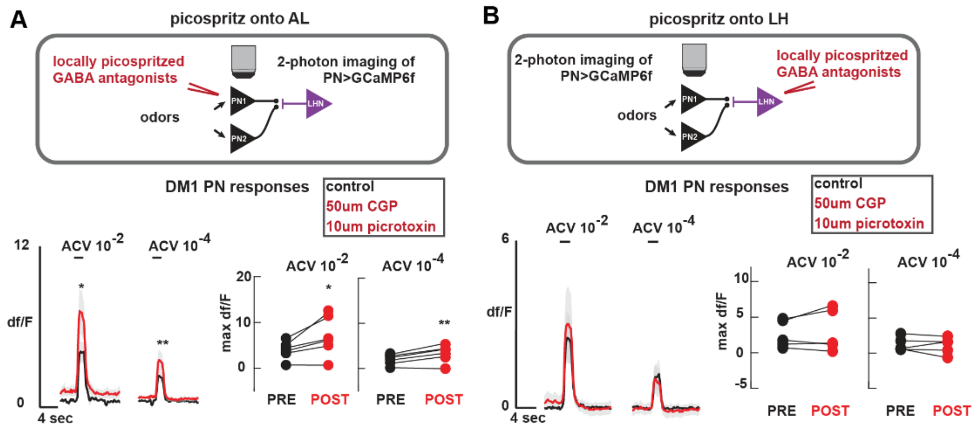


**Figure S4. Local GABA antagonism remains within the lateral horn.**

**(A)** Top: schematic of local GABA receptor antagonism in the antennal lobe while imaging in the antennal lobe. Bottom: Picrotoxin (10um) and CGP54626 (50μM) were pressure ejected into the antennal lobe. Pre- and postdrug odor-evoked GCaMP6f activity were compared within the same flies in the DM1 glomerulus (n=6). For ACV at $10^{-2}$× and $10^{-4}$× concentrations, GABA receptor blockade increased odor-response magnitude (paired t test, p = 0.0365 ACV $10^{-2}$, p = 0.0103, ACV $10^{-4}$).

**(B)** Same as (A), but with local GABA receptor antagonism in the lateral horn while imaging in the antennal lobe. Odor-response magnitude was unchanged in pre vs. post drug conditions (paired t test, p = 0.4090 ACV $10^{-2}$, p = 0.6039, ACV $10^{-4}$). Applying drugs to the lateral horn therefore does not alter odor coding in the antennal lobe.

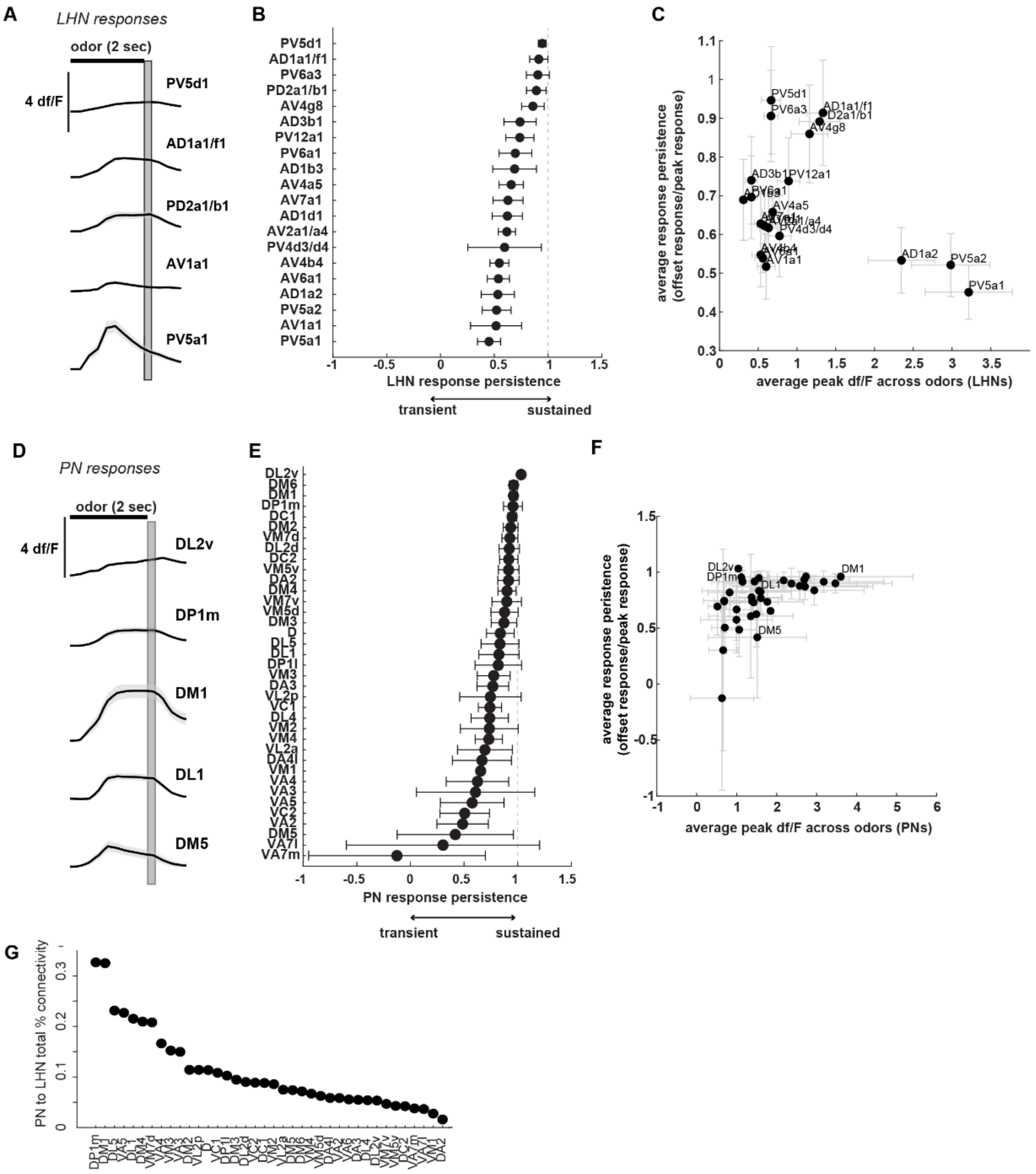


**Figure S5. LHNs have diverse temporal dynamics and receive largest synaptic input from sustained PNs. Adaptation dynamics are similar across different lengths of odor stimulation.**

**(A)** Temporal response profiles for 5 LHN types averaged across all 12 odors (shading is s.e.m.). Some LHNs sustain responses during odor delivery (e.g., PV5d1). Others adapt quickly even during ongoing odor delivery (e.g., PV5a1). Adaptation ratio is calculated as dF/F at the end of odor stimulation divided by df/F during the initial peak. The window used for the end of odor stimulation is denoted by the grey vertical bar.

**(B)** Different LHN types have diverse adaptation ratios (one-way ANOVA, $p = 3.406e^{-21}$)

**(C)** Relationship between LHN response persistence and average response magnitude (error bars are s.e.m. across odors). LHNs with the largest odor responses can be transient (e.g., PV5a1) or sustained (e.g., PD2a1/b1), suggesting that response persistence does not have a one-to-one correspondence with response magnitude.

**(D)** PN average temporal response profiles across odors (shaded region is s.e.m.). Some PNs exhibit sustained responses (e.g., DP1m, DL2v). Others adapt quickly before the end of odor stimulation (e.g., DM5).

**(E)** Different PN types have diverse response persistence (one-way ANOVA, $p = 2.838^{-16}$).

**(F)** Relationship between PN response persistence and average response magnitude (error bars are s.e.m. across odors). PNs with sustained dynamics (e.g., DP1m, DL2v, and DM1) vary in absolute response magnitude, suggesting that response persistence does not have a one-to-one correspondence with response magnitude.

**(G)** LHNs in our dataset receive the highest synaptic input from PNs with sustained temporal dynamics (DM1, DP1m). Therefore, while PNs do exhibit diverse temporal profiles, most PNs providing synaptic input to LHNs in our sample are sustained.
